# Retroviral insertions contributed to the divergence of human and chimpanzee brains

**DOI:** 10.64898/2025.12.12.693858

**Authors:** Patricia Gerdes, Ofelia Karlsson, Raquel Garza, Diahann A.M. Atacho, PingHsun Hsieh, Bharat Prajapati, Chandramouli Muralidharan, Anita Adami, Carrie Davis-Hansson, Emily M. Johansson, Laura Castilla-Vallmanya, Wiktor West, Meghna Vinoud, Lucian Domitrovic, Pia A. Johansson, Jenny Johansson, Christopher H. Douse, Chandrasekhar Kanduri, Patric Jern, Evan E. Eichler, Johan Jakobsson

**Affiliations:** Laboratory of Molecular Neurogenetics, Department of Experimental Medical Science, Wallenberg Neuroscience Center and Lund Stem Cell Center, BMC A11, Lund University, 221 84 Lund, Sweden; Department of Genetics, Cell Biology, and Development, Institute for Health Informatics and Bioinformatics and Computational Biology Graduate Program, University of Minnesota, Twin Cities, MN, USA; Department of Medical Biochemistry and Cell Biology, Institute of Biomedicine, University of Gothenburg, SE-40530 Gothenburg, Sweden; Laboratory of Epigenetics and Chromatin Dynamics, Department of Experimental Medical Science, Wallenberg Neuroscience Center and Lund Stem Cell Center, BMC A11, Lund University, 221 84 Lund, Sweden; Department of Medical Biochemistry and Microbiology, Uppsala University, Uppsala, Sweden; Department of Genome Sciences, University of Washington School of Medicine, Seattle, WA, USA; Howard Hughes Medical Institute, University of Washington, Seattle, WA, USA

## Abstract

Over the past 5–7 million years, humans and chimpanzees have diverged in brain size, structural complexity, and cognitive abilities despite high conservation of protein-coding genes. Notably, the endogenization and proliferation of retroviral infections within host genomes has introduced numerous species-specific regulatory elements that have the potential to influence gene regulation. However, the role of these endogenous retroviruses in hominoid brain evolution remains unclear. A burst of lineage-specific PTERV1 retroviruses recently invaded the chimpanzee genome but are absent in humans. We conducted an epigenomic analysis of PTERV1 insertions in chimpanzee neural organoids and found that they are heavily covered by DNA methylation, representing more than 150 species-specific heterochromatin domains with the capacity to influence gene regulatory networks. We identified one such chimpanzee-specific PTERV1 insertion on chromosome 19 that blocks the expression of the long noncoding RNA LINC00662, via DNA methylation spread to the adjacent genomic region. The expression of LINC00662 was restored in chimpanzee induced pluripotent stem cells when we deleted the PTERV1 insertion using CRISPR editing. We found that LINC00662, a human-specific RNA, is highly expressed in the developing brain and plays an important role in the posttranscriptional control of neuronal maturation, axon outgrowth, and neural organoid development. In summary, our findings describe how endogenous retroviral insertions contributed to the functional divergence of the human and chimpanzee brains. This provides a new mechanism by which retroviral pandemics influenced primate brain speciation.

## Introduction

Among primates, the human brain stands out as the largest and most complex due in part to an increase in cortical neurons and increase in synaptic connectivity. With a size roughly three times that of our closest living evolutionary relative, the chimpanzee, it possesses unique cognitive capabilities^1^. The underlying genetic divergences responsible for these differences are still poorly understood^2^. Humans share more than 96% of protein-coding sequences with chimpanzees, making it unlikely that species-specific protein-coding variants are the sole drivers of differences in brain size and complexity^3^. Rather, a substantial part of the genetic basis for the differences between nonhuman primate and human brains is likely to lie in the noncoding part of the genome, either as noncoding single base-pair substitutions or as structural variants^4–9^. Previous studies have implicated transposable element (TE)-driven remodeling of gene regulatory networks in shaping human-specific gene expression patterns during brain development and evolution^5,10–15^. However, the contributions of TEs to primate brain speciation remain largely unexplored.

A large proportion of primate genome sequences is made up of TEs, with one major family being endogenous retroviruses (ERVs), which have expanded throughout primate evolution. In humans and chimpanzees, at least 8% of the genome consists of ERVs, including both full-length proviruses and solo long terminal repeat (LTR) fragments^9,16,17^. Most primate ERVs are believed to have originated from exogenous retroviral integration events in the germline. These ERVs then expanded through reinfection or intercellular retrotransposition, and were subsequently fixed in the genome, probably through population bottlenecks^18^. Comparative analyses of closely related primate genomes have revealed substantial differences in ERV composition between species which provides a genomic fossil of ancient retroviral pandemics. These differences demonstrate that ERVs represent a rich source of species-specific structural variants^9,19,20^. Furthermore, ERVs have the potential to impact genomes by altering gene regulatory networks. For example, ERVs harbor promoter and enhancer sequences that can affect the expression patterns of nearby genes *in cis*^21,22^. In addition, ERVs are preferential sites for DNA methylation, which may influence the epigenetic status of their integration sites and silence genes in the vicinity^23–26^. This capacity of ERVs to influence host gene expression has led to the idea that ERVs may be actively involved in speciation by contributing to species-specific gene regulatory networks^27–30^.

The rate of ERV insertions into the human genome has decreased during the most recent phase of human evolution. After the chimpanzee-hominoid split about 5-7 million years ago, only a small number of HERV-K elements integrated and became fixed in the human genome^31–33^. However, this lack of a major burst of ERV integrations into the genome is unique for humans among African great apes^34^. The chimpanzee genome was recently invaded by a retrovirus family, *Pan troglodytes* endogenous retrovirus 1 (PTERV1), which is completely absent from the human genome. PTERV1 generated at least 150 fixed insertions, making it one of the most abundant ERV families in the chimpanzee genome^9,35,36^. These PTERV1 insertions are thought to have originated from infections that occurred in the ancestors of African apes about 5-7 million years ago^36^. PTERV1 elements are found in chimpanzees, bonobos and gorillas; however, only a single orthologous insertion has been identified between chimpanzee and gorilla lineages, suggesting independent expansion in these species^9^ (**Fig. 1A**). Notably, the absence of PTERV1 insertions from the human genome suggests that either the human and chimpanzee lineages did not occupy the same habitat at the time or that germline-competent cells of the human ancestor were resistant to this infection. In line with this latter idea, each primate species encodes a TRIM5α protein with different antiviral specificity and it has been shown that the human version of TRIM5α prevents infection of human cells with PTERV1^37^.

**Figure 1.**
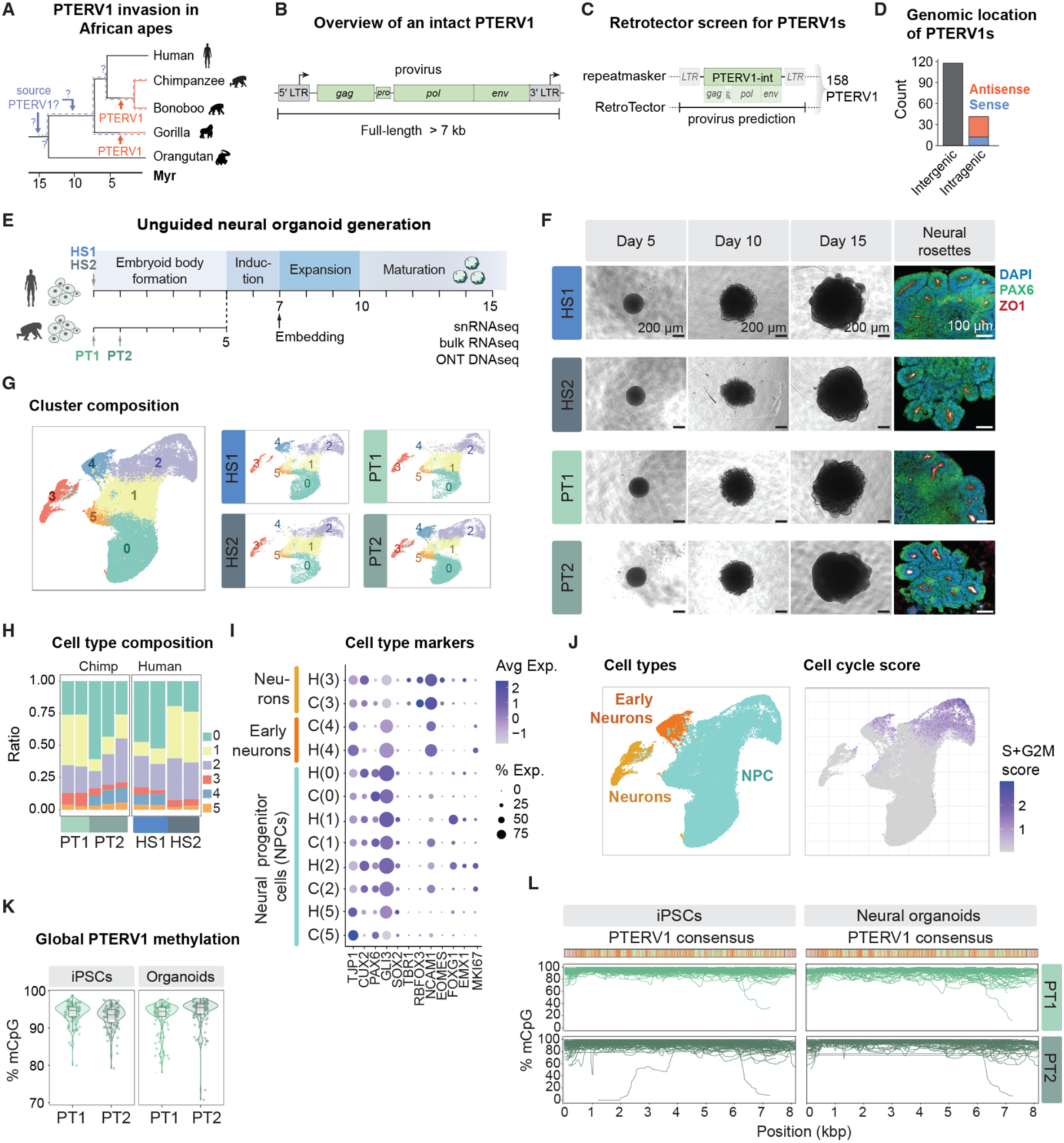
**A** Phylogenetic tree depicting the estimated timepoints of PTERV1 infection and colonization of Great Ape genomes. **B** Structure of a PTERV1 element. A typical full-length PTERV1 is about 7 kb in length and consists of two 400-600 bp long LTRs (grey boxes) flanking the retroviral genes (*gag, pro, pol* and *env*; green boxes). The LTRs have promoter function (black arrows). **C** Schematic of Retrotector screen and analysis of PTERV1 proviruses in the chimpanzee reference genome (CLINT_PTRv2/panTro6). **D** PTERV1 integration sites in the chimpanzee reference genome. Integration sites (inter-versus intragenic) were defined based on the panTro6 RefSeq transcript annotation. PTERV1 orientation was defined based on 5′to 3′ synthesis and classified with respect to sense/antisense orientation of the RefSeq transcript annotation. **E** Schematic of timeline for unguided neural organoid generation from human and chimpanzee iPSCs. Different colors show changes in media and stages of the protocol. Note that PT2 organoids have an EB stage that is 1 day shorter than the other cell lines. **F** *Left:* Bright field images of human (HS1 and HS2) and chimpanzee (PT1 and PT2) neural organoids at day 5, 10 and 15 of differentiation. Scale bar, 200 um. *Right:* Representative immunofluorescence images of neural rosettes at day 15 of neural organoid differentiation with staining for neural identity marker PAX6 (green) and tight junction marker ZO1 (red). DAPI is shown in blue. Scale bar, 100 μm. **G** UMAPs coloured by nuclei clusters found in day 15 unguided neural organoids with all nuclei (left) and split per cell line (right). **H** Barplot showing cluster composition per sample. **I** Expression of selected cellular markers across species. Color showing average expression in cluster, dot size showing percentage of nuclei in cluster where the gene was detected. **J** *Left*: UMAP coloured by cell types. *Right*: UMAP coloured by cell cycle score (sum of gene module scores of genes associated to S-phase and G2M-phase. Seurat::CellCycleScoring). **K** PTERV1 CpG methylation ascertained by ONT DNA sequencing of chimpanzee iPSCs and day 15 unguided neural organoids. Dots represent single PTERV1 loci. **L** Composite FL-PTERV1 methylation profiles. A schematic of the FL-PTERV1 consensus is shown at the top with CpG dinucleotides indicated in orange.

The endoretroviral pandemic underlying the integration of PTERV1 sequences into the ancestral chimpanzee genome is likely to have had a major negative impact on the fitness and size of the population at that time and could have contributed to the speciation between the human and chimpanzee lineages by changing population structure and habitat. In addition, a more indirect but long-lasting effect of this retroviral pandemic could be that fixed PTERV1 insertions affected gene regulatory networks, thereby acting as a source of genetic divergence that could contribute to the speciation between human and chimpanzee ancestors. However, experimental evidence for this hypothesis is so far lacking.

In this study, we conducted a global epigenomic analysis of PTERV1 insertions in chimpanzee neural organoids. We found that these elements are heavily covered by DNA methylation, representing species-specific heterochromatin domains. We identified a PTERV1 insertion on chimpanzee chromosome 19 that affects gene regulatory networks in brain development through the transcriptional repression of the long noncoding RNA (lncRNA) LINC00662, via accumulation of DNA methylation. We were able to restore LINC00662 expression in chimpanzee cells using CRISPR-based deletion of this PTERV1 in chimpanzee induced pluripotent stem cells (iPSCs). We found that LINC00662 is a human-specific transcript with high expression in the developing brain. It is localized in the cytoplasm, where it plays a key role in the maturation of neurons, thereby influencing the development of neural organoids. In summary, this study describes how retroviral insertions contributed to the functional divergence of the human and chimpanzee brain, providing a new mechanism by which retroviral infections influenced primate brain speciation.

## Results

### PTERV1s are covered by DNA methylation in chimpanzee iPSCs and neural organoids

A full-length PTERV1 is typically 7-10 kb in length and encodes the retroviral genes *gag, pro pol,* and *env*, which are flanked by two long terminal repeats (LTRs) that are identical at the time of integration and usually 400-600 bp long^35,36^ (**Fig. 1B**). Due to the fragmented RepeatMasker annotation of ERVs in reference genomes, which splits ERVs into LTRs and internal regions, we used the RetroTector software^38^ to screen the chimpanzee reference genome (CLINT_PTRv2/panTro6) for PTERV1 proviruses. We found 158 PTERV1 elements which ranged in size from 2 to 14 kb with an average length of 7.5 kb (**Fig. 1C**, **Table S1**). In agreement with previous analysis^9^, we found that the majority of these PTERVs were intergenic and integrated preferentially in antisense direction when located within genes, indicating purifying selection (**Fig. 1D**).

Comparative evolutionary analyses of hominoid brain development have long been hampered by the lack of availability of chimpanzee tissue from early developmental periods. However, with the generation of iPSCs of chimpanzee origin, it has now become possible to develop *in vitro* cell culture models to study hominoid brain development^39–43^. To investigate the epigenetic status of individual PTERV1 elements and their potential effect on gene regulatory networks in brain development, we optimized a protocol to generate chimpanzee and human unguided neural organoids under near-identical conditions to obtain highly comparable organoids for the two species (**Fig. 1E**). We observed similar size and morphology between chimpanzee and human organoids up to 15 days of differentiation using iPSCs from two individuals of each species (**Fig. S1A**). At this developmental stage, organoids from both species showed similar neural rosette formation typical for unguided neural organoids, as monitored with immunohistochemical staining for PAX6, a neural identity marker, and ZO1, a tight junction protein^44,45^ (**Fig. 1F**). Single nuclei RNA-seq analysis (snRNA-seq) confirmed a similar cell type composition of the chimpanzee and human organoids, with most cells being neural progenitor cells (NPCs) or early-born neurons at day 15 of neural organoid development (**Fig. 1G-I**).

To investigate the epigenetic status of PTERV1s in chimpanzee iPSCs and unguided neural organoids, we performed genome-wide CpG DNA methylation profiling using long-read DNA sequencing from Oxford Nanopore Technologies (ONT), as this approach allows the analysis of DNA methylation patterns of individual PTERV1 loci at single base-pair resolution. This analysis revealed that almost all PTERV1 loci in iPSCs and day 15 unguided neural organoids are completely methylated (**Fig. 1J-K**). Since PTERV1 integrations are absent from the human genome, this result demonstrates that the PTERV1 insertions represent 158 chimpanzee-specific local heterochromatin domains.

### The expression of LINC00662 in chimpanzee is silenced by a PTERV1 insertion

We hypothesized that methylated PTERV1 insertions might affect gene expression in *cis* through the spread of local heterochromatin. Therefore, we intersected PTERV1 insertions with genes that have a homologue in both species (**Fig. 2A**). Among the 46 genes that intersected with a PTERV1 insertion, we found 11 genes, including both protein-coding and non-coding transcripts, showed differential expression when comparing bulk RNA-seq data from human and chimpanzee unguided neural organoids (|log2FoldChange| > 1; padj < 0.05 Wald test, DESeq2) (**Fig. 2B**). The top candidate of these genes was LINC00662, a lncRNA which is highly expressed in human unguided neural organoids but completely silenced in chimpanzee organoids (**Fig. 2B**).

**Figure 2.**
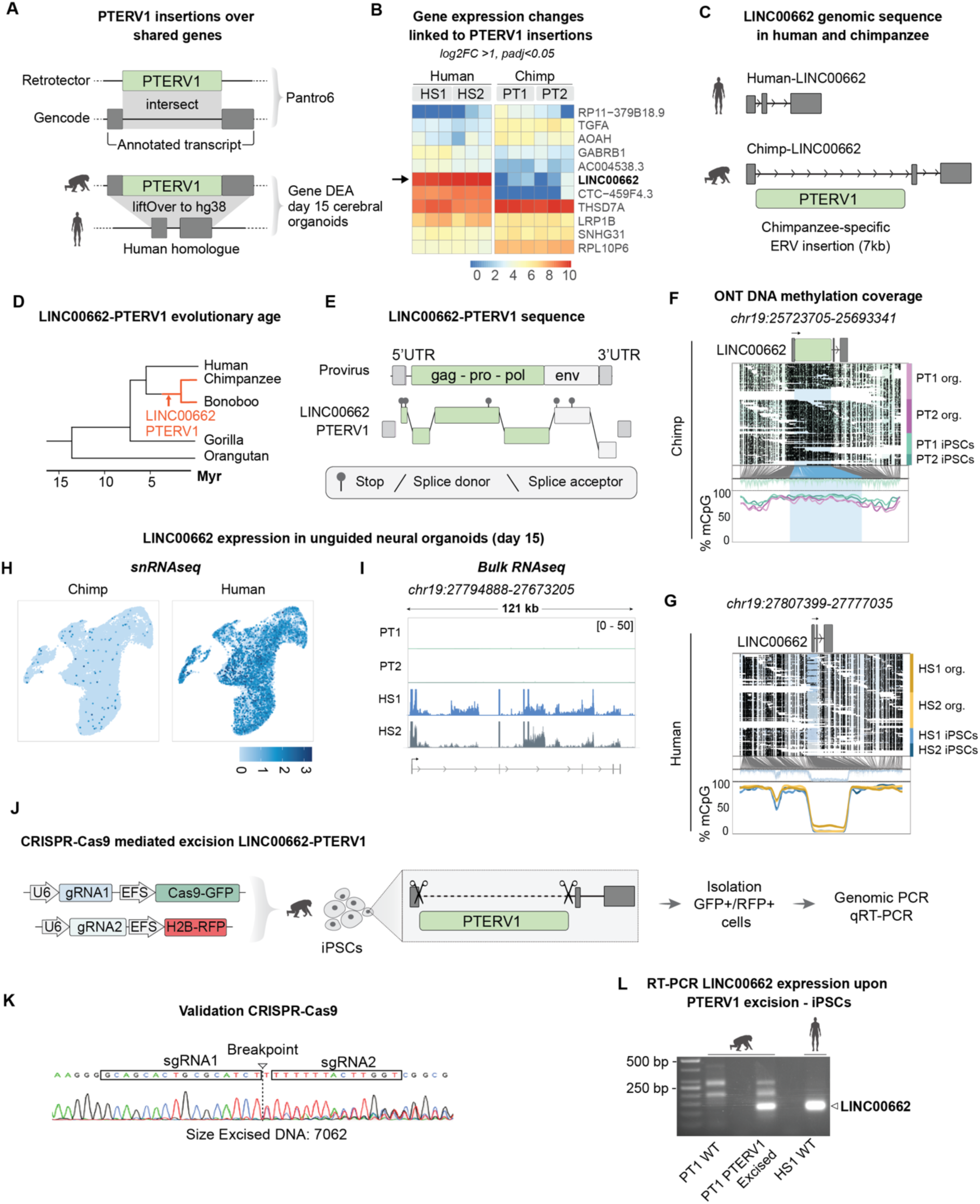
**A** Workflow for comparing PTERV1 containing transcripts and their human homologues expression in day 15 unguided neural organoids. **B** Expression of transcripts with a PTERV1 insertion in the chimpanzee genome in day 15 human HS1 (n=3) and HS2 (n=3) and chimpanzee PT1 (n=3) and PT2 (n=3) unguided neural organoids as determined by bulk RNA sequencing. Results are presented as LFC >1 and padj <0.05. **C** Schematic of the human and chimpanzee *LINC00662* sequence with indicated location of a PTERV1-insertion in the chimpanzee genome. **D** Phylogenetic tree with indication of the estimated evolutionary age of *LINC00662*-PTERV1. **E** Schematic of the PTERV1 provirus and three frame chain view of LINC00662-PTERV1 as determined by RetroTector showing stop codons and frameshifts. **F** DNA methylation profile of the *LINC006622*-PTERV1 locus obtained by ONT DNA sequencing. The first top panel shows a PTERV1 insertion in sense to intron 1 of *LINC00662* in chimpanzee. The second panel displays ONT read alignments from iPSC lines PT1 and PT2 and iPSC-derived unguided neural organoids. Unmethylated CpGs colored in color of respective sample, methylated CpGs colored in black. The third panel indicates the CpG position including those corresponding to the PTERV1 (shaded in blue). The fourth panel shows fraction of methylated CpGs. **G** As in **F** but for human *LINC006622* locus (PTERV1 not present). Samples are human iPSC lines HS1 and HS2, human iPSC-derived unguided neural organoids and human fetal forebrain samples (n = 2). **H** Expression of LINC00662 in human (HS1 and HS2) iPSC-derived unguided neural organoids and chimpanzee (PT1 and PT2) iPSC-derived organoids at day 15 according to snRNA-seq. **I** Snapshot of genome browser tracks showing expression of LINC00662 in human (HS1 and HS2) and chimpanzee (PT1 and PT2) iPSC derived unguided neural organoids at day 15 according to bulk RNA sequencing. **J** Workflow for CRISPR-Cas9-mediated excision of the PTERV1-insertion in chimpanzee iPSC (PT1). **K** Chromatogram showing sequencing results upon CRISPR-Cas9 mediated excision of *LINC00662-*PTERV1 in PT1. **L** RT-PCR product of chimpanzee iPSC, chimpanzee PTERV1-excised iPSCs and human iPSC size separated on an agarose gel. The expected amplicon size of human *LINC00662* and chimpanzee PTERV-1 excised *LINC00662* are indicated with black arrow.

Pairwise sequence alignment of the human and chimpanzee *LINC00662* genome sequences revealed that the chimpanzee locus contains a 6915 nt PTERV1 insertion integrated into the negative strand in sense within the first intron of *LINC00662* **(Fig. 2C)**. We found no other major differences between the human and chimpanzee genome at this locus (**Fig. S2A**), indicating that the PTERV1 insertion is likely to be the underlying cause for the transcriptional differences. Using multiple sequence alignments, we found that the *LINC00662*-PTERV1 insertion is present in the chimpanzee and bonobo genomes, but absent in the human and orangutan genomes. This suggests that the insertion occurred between the splits of the human and chimpanzee lineages, and the chimpanzee and bonobo lineages, which occurred between 1.5 and 5 million years ago **(Fig. 2D, Fig. S2B)**. Analysis of the 5′ and 3′ LTR divergence of the flanking LTR elements of the *LINC00662*-PTERV1 showed a low divergence (1.2% mutation rate), confirming the young evolutionary age of this retroviral insertion **(Fig. 2D)**. Structural analysis of the *LINC00662*-PTERV1 provirus sequence, using the RetroTector software, confirmed that it is of gamma-retrovirus origin and contains the retroviral genes *gag, pro, pol* and *env* (**Fig. 2E**). The gamma-retrovirus gag-pro-pol polyprotein is normally expressed in a continuous reading frame with a read-through stop after gag. The *LINC00662*-PTERV1 provirus gag-pro-pol contains multiple inactivating frameshifts and stop codons. In addition, the *LINC00662*-PTERV1 env displays frameshifts and stop codons **(Fig. 2E)**. In summary, the *LINC00662*-PTERV1 sequence diverges only slightly from the PTERV1 consensus sequence, but it is likely replication-deficient due to frameshifts and premature stop codons.

As PTERV1 insertions are highly methylated (**Fig. 1G**), we hypothesized that the *LINC00662*-PTERV1 silences the *LINC00662* promoter by introducing a local heterochromatin domain. To investigate the epigenetic status of the *LINC00662* locus in humans and chimpanzees we used the ONT DNA sequencing data from iPSCs and neural organoids. In chimpanzee iPSCs and neural organoids, the *LINC00662-*PTERV1 insertion was highly methylated, as were the flanking genomic regions **(Fig. 2F)**. On the contrary, we found low methylation coverage across the *LINC00662* locus in human iPSCs and neural organoids **(Fig. 2G)**. These results suggest that the PTERV1 insertion in the chimpanzee *LINC00662* locus attracts DNA methylation that spreads into the flanking genome, contributing to the differences in DNA methylation patterns at this locus between humans and chimpanzees.

To further dissect the transcriptional divergence of *LINC00662*, we used our neural organoid bulk RNA-seq and snRNA-seq data, which confirmed that LINC00662 was exclusively expressed in human samples **(Fig. 2H-I)**. LINC00662 was expressed throughout the different cell populations in unguided neural organoids, ruling out the possibility that the human-specific expression was due to a specific subpopulation in the human samples. We used our previously published CUT&RUN epigenomic profiling data of human and chimpanzee forebrain NPCs grown in 2D^6^, which showed a striking human-specific enrichment of the transcription-related epigenetic mark H3K4me3 over the promoter of *LINC00662*, confirming that the observed differences in RNA levels were due to differences in transcriptional activity **(Fig. S2C)**. To further verify the human-specific expression of LINC0066*2* we analyzed publicly available RNA sequencing data of human, chimpanzee, and rhesus macaque cerebral organoids as well as of human, chimpanzee, and bonobo NPCs^46,47^. These data sets revealed a high expression of LINC00662 in human, but not chimpanzee, rhesus macaque or bonobo neuronal cell models **(Fig. S2D-E).** To further verify that our results were not limited to the 2 chimpanzee individuals used to derive the iPSCs in this study, a dataset of iPSCs of 8 additional individuals confirmed human-specific expression of LINC00662^48^ **(Fig. S2F)**.

To investigate mechanistically whether the PTERV1 insertion is responsible for the transcriptional silencing of *LINC00662*, we used CRISPR-Cas9 gene editing to excise it in chimpanzee iPSCs. Two guide RNAs were designed to target the flanking regions of the *LINC00662*-PTERV1 and cloned into lentiviral vectors that also express Cas9 (**Fig. 2J**). Amplicon sequencing of chimpanzee iPSCs transduced with the CRISPR-Cas9 vector confirmed the excision of the ∼7kb PTERV1 (**Fig. 2K**). RT-PCR analysis showed that upon excision of PTERV1, LINC00662 was expressed in chimpanzee iPSCs (**Fig. 2L**). Thus, this experiment demonstrates that the *LINC00662*-PTERV1 insertion mediates transcriptional silencing of *LINC00662* in chimpanzees as PTERV1 excision restores LINC00662 expression in chimpanzee cells.

### LINC00662 is a highly abundant trans-acting lncRNA expressed during human brain development

To investigate LINC00662 expression during human brain development *in vivo*, we analyzed publicly available bulk RNA sequencing data from different stages of the developing and adult human brain^11^. We found that LINC00662 expression peaks in early brain development at around 10-12 weeks post conception (PC), followed by gradual downregulation in adulthood (**Fig. 3A**). snRNA-seq analysis of human fetal forebrain development using our previously published dataset^49^ showed that LINC00662 is expressed throughout the developing human forebrain at weeks 7- and 11-weeks PC (**Fig. 3B**). We further confirmed that LINC00662 is expressed in human fetal forebrain development at 7- and 11-weeks PC using our previously published bulk RNA-seq and H3K4me3 CUT&RUN dataset^49^. Additionally, ONT direct long-read RNA sequencing showed that LINC00662 expression in the fetal human brain at 9- and 10-weeks PC starts at the TSS, which overlaps with the H3K4me3 peak, and that it is a spliced lncRNA that exists in several different isoforms due to alternative transcription stop sites (**Fig. 3C**). ONT long-read DNA sequencing revealed low DNA methylation coverage across the *LINC00662* locus in human fetal brain tissue at weeks 7 and 11 PC (**Fig. 3D).** Taken together, these results demonstrate that LINC00662 is a highly abundant noncoding transcript expressed during early human brain development.

**Figure 3.**
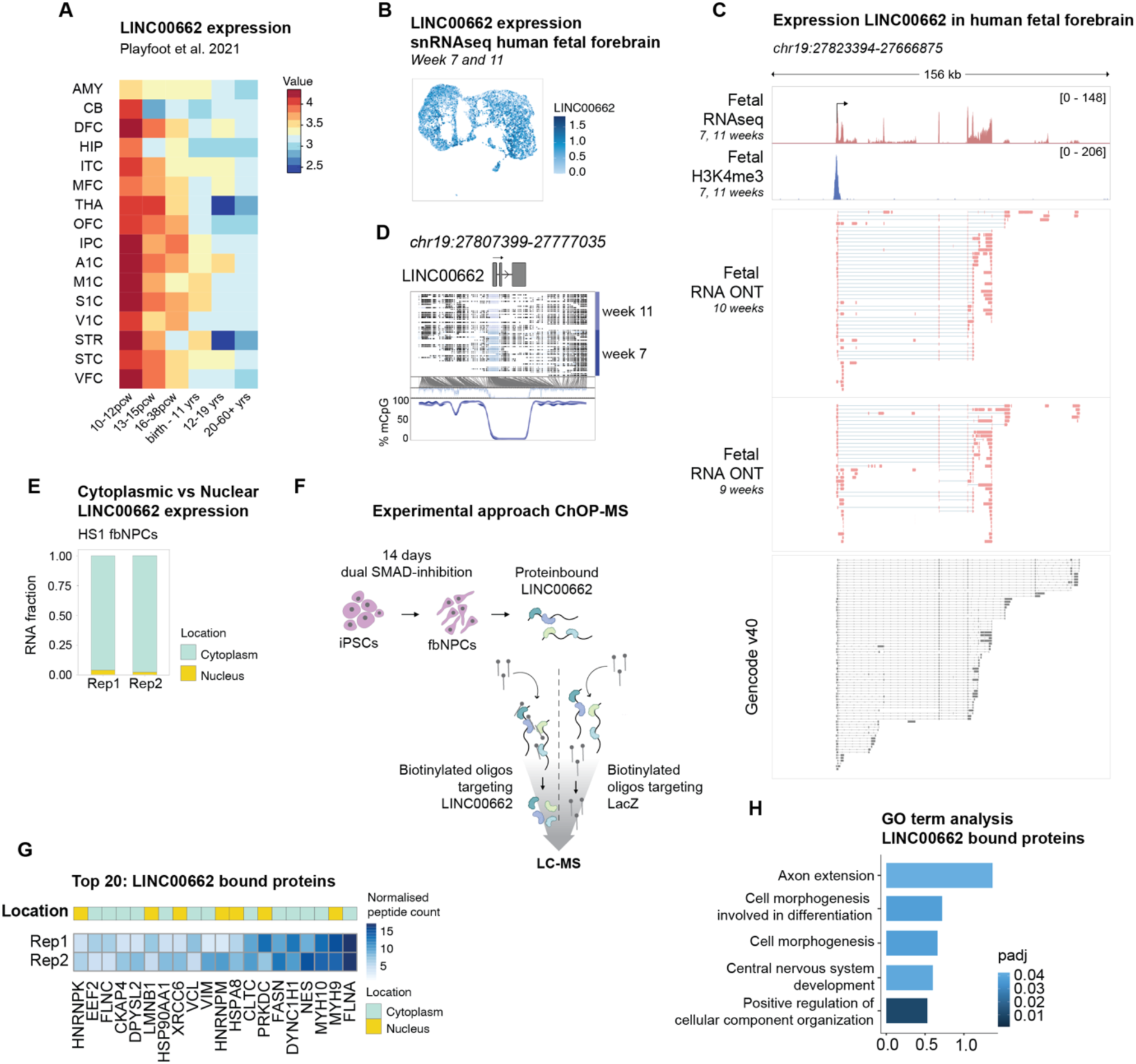
**A** Bulk RNA-seq showing LINC00662 expression in human fetal and adult brain samples presented as Log_2_ CPM. Data from Playfoot et al 2021^11^. AMY (amygdala), CB (cerebellar cortex), DFC (dorsolateral prefrontal cortex), HIP (hippocampus), ITC (inferior temporal cortex), MFC (medial prefrontal cortex), THA (mediodorsal nucleus of the thalamus), OFC (orbital prefrontal cortex), IPC (posterior inferior parietal cortex), A1C (primary auditory (A1) cortex), M1C (primary motor (M1) cortex), S1C (primary somatosensory (S1) cortex), V1C (primary visual (V1) cortex), STR (striatum), STC (superior temporal cortex), VFC (ventrolateral prefrontal cortex). **B** Expression levels of LINC00662 in human fetal forebrain samples at different gestational ages. **C** Snapshot of genome browser tracks showing LINC00662 expression in human fetal forebrain samples as determined by bulk RNA sequencing (top panel), CUT&RUN analysis of H3K4me3 (middle panel) and reads from direct RNA ONT (bottom panel). **D** DNA methylation profile of the *LINC006622* locus obtained by ONT sequencing of two fetal forebrain samples. The first top panel shows *LINC00662* in the human genome. The second panel displays ONT read alignments. Unmethylated CpGs colored in color of respective sample, methylated CpGs colored in black. The third panel indicates the CpG position. The fourth panel shows fraction of methylated CpGs in the region. **E** Subcellular localization of LINC00662 in human HS1 fbNPCs as determined by qRT-PCR analysis of subsequent cytosolic and nuclear RNA extractions (n= 2, Rep1 and Rep2). **F** Illustration showing workflow for ChOP mass spectrometry followed by characterization using liquid chromatography-mass spectrometry of proteins bound to LINC00662 in human fbNPCs. **G** Heatmap showing normalized values of top 20 proteins bound to *LINC00662* and their cellular localization in human fbNPCs. (n=2, Rep1 and Rep2). **H** Gene Onthology (GO) term analysis of proteins bound to LINC00662. Results were generated using StringDb v11.5.

LncRNAs act *in cis* by influencing the expression or chromatin state of nearby genes or *in trans* by executing functions throughout the cell, for example in the cytoplasm^50^. To investigate the cellular localization of LINC00662, we performed a cellular fractionation assay in iPSC-derived human fbNPCs, which revealed an almost exclusive presence of LINC00662 transcripts in the cytoplasmic fraction (**Fig. 3E**), indicating that LINC00662 acts in *trans*. Since many lncRNAs act as protein-binding RNAs, we determined the protein interactome of LINC00662 using chromatin oligo affinity purification followed by shot-gun proteomics (ChOP-MS). Using targeted biotinylated oligonucleotides, we pulled down proteins bound by LINC00662, which we characterized using liquid chromatography-mass spectrometry (LC-MS/MS) (**Fig. 3F**). As a negative control we used oligonucleotides targeting *lacZ*, representing a sequence not found in the human genome. The ChOP-MS results confirmed that LINC00662 mainly interacts with proteins found in the cytoplasm, further indicating that this lncRNA acts *in trans* (**Fig. 3G**). Gene ontology (GO) term analysis of the LINC00662-bound proteins indicated an enrichment of terms related to axon extension and neural development (**Fig. 3G-H**). For example, proteins that are in complex with LINC00662 include FLNA, MYH10, NES, VIM, and DPYSL2. These proteins are all related to the cytoskeleton and are well-documented examples of proteins that are linked to neurite extension and brain development^51–58^ (**Fig. 3G**). Taken together, these results suggest that LINC00662 may play an important role in the cytoplasm in neural progenitors by interacting with proteins associated with the cytoskeleton and neurite extension.

### LINC00662 plays a role in neurite extension in neural organoids

The LINC00662-protein interactome analysis indicated a role in neurite extension during brain development. To investigate the functional relevance of LINC00662, we developed a CRISPRi strategy to silence its expression in neural organoids. We designed two guide RNAs (gRNAs) to target unique genomic locations in the vicinity of the TSS and co-expressed these with a Krüppel-associated box (KRAB) transcriptional repressor domain fused to catalytically dead Cas9 (KRAB-dCas9) (**Fig. 4A**). Lentiviral transduction of human iPSCs resulted in efficient, almost complete silencing of LINC00662 in iPSCs using both gRNAs (**Fig. 4B, Fig. S3A**). To investigate whether LINC00662 is involved in the neural differentiation and neurite extension, we generated *LINC00662-*CRISPRi unguided neural organoids. For this experiment, we grew organoids for 15 and 30 days to capture the period when newborn neurons begin to mature and form their first neurites.

**Figure 4.**
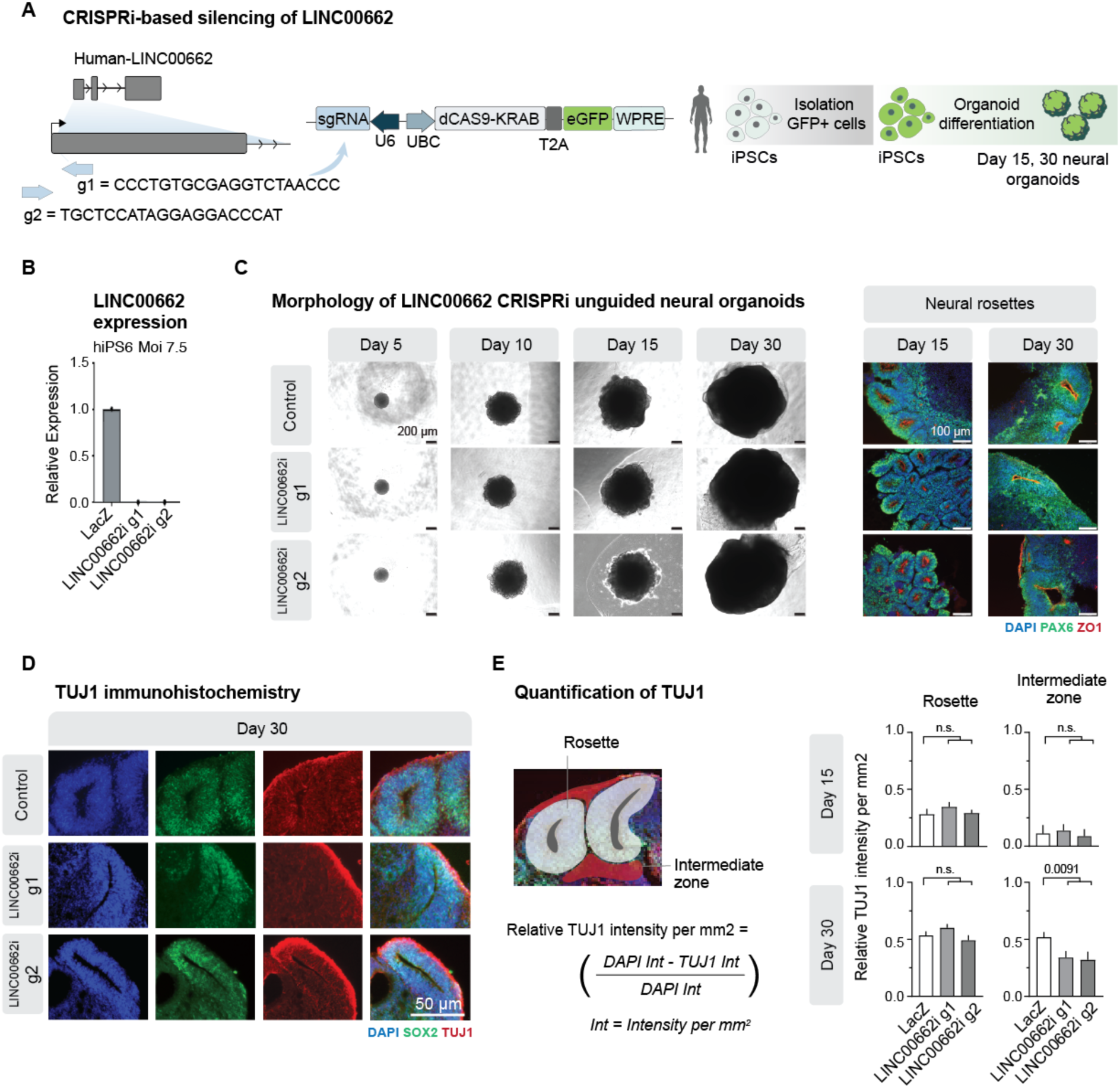
**A** Schematics of experimental design for *LINC0062*-CRISPRi in human iPSC-derived unguided neural organoids. **B** qRT-PCR analysis of relative LINC00662 expression in human iPSC: control (LacZ) and *LINC00662*-CRISPRi (gRNA1, gRNA2). **C** Left: Brightfield images of iPSC-derived *LINC00662*-CRISPRi unguided neural organoids. Black scale bars = 200 μm. Right: Immunohistochemistry of day 15 and day 30 *LINC00662*-CRISPRi unguided neural organoids of PAX6 (green), ZO-1 (red) and DAPI (blue). White scale bars = 100 μm. **D** Immunohistochemistry of day 30 *LINC00662*-CRISPRi unguided neural organoids of SOX2 (green), Tuj1 (red) and DAPI (blue). White scale bars = 50 μm. **E** Left: Illustration of regions of interest for analysis. Right: Quantification of the normalized mean Tuj1 intensities in the different regions of interest, 2-3 rosettes and intermediate zones per organoid from different planes, from six organoids per condition derived from two independent batches from day 15 and day 30 *LINC00662*-CRISPRi organoids. Two-tailed Student’s *t* test.

We found that *LINC00662*-CRISPRi silencing did not affect organoid formation or growth, and the resulting organoids displayed characteristic neural rosettes after 15 days of growth, as visualized by PAX6/ZO1 staining (**Fig. 4C**). To visualize the process of neuronal differentiation, we stained for TUJ1 (beta-III-tubulin), which labels the microtubule cytoskeleton in the soma and neurites of newly committed neuroblasts. At day 30 we observed extensive TUJ1 labeling in the control organoids, whereas the intensity of TUJ1 staining was markedly lower in the *LINC00662*-CRISPRi organoids (**Fig. 4D**). We quantified TUJ1 staining intensity in the neural rosettes, which are composed of progenitor cells, and the intermediate zone where committed neuroblasts and their neurite extensions are located. As expected, we detected low levels of TUJ1 staining in both the neural rosettes and the intermediate zone on day 15, with no difference between the control and *LINC00662*-CRISPRi organoids (**Fig. 4E**, 2-3 rosettes and intermediate zones per organoid from different planes, from six organoids per condition derived from two independent batches from day 15 and day 30). At day 30, we observed a clear increase in TUJ1 labeling intensity in the intermediate zone of control organoids compare to the day 15 timepoint, confirming the formation of committed neuroblasts with neurite extensions at this time point (**Fig. 4E**). Interestingly, TUJ1 intensity was significantly reduced in the intermediate zone, but not in the neural rosettes, in *LINC00662*-CRISPRi neural organoids (both gRNAs) (**Fig. 4E**). This data suggests that LINC00662 is directly involved in the formation of the neurites of newly committed human neuroblasts.

### Transcriptional alterations in LINC00662-CRISPRi neural organoids

Because LINC00662 is primarily found in the cytoplasm, where it binds to proteins associated with the cytoskeleton and neurite extension, this lncRNA likely regulates neurite extension through a post-transcriptional mechanism. To investigate if silencing of *LINC00662* has a downstream impact on cell state and transcription we performed snRNA-seq analysis of *LINC00662*-CRISPRi and control neural organoids at 15 and 30 days of differentiation. High-quality data were generated from a total of 119674 nuclei, including 86098 from *LINC00662*-CRISPRi organoids (two gRNAs, in total 61 organoids) and 33576 from control organoids (*lacZ*-gRNA, in total 24 organoids). We performed an unbiased clustering analysis to identify and quantify the different cell types present in the organoids. Seven separate clusters were identified (**Fig. 5A-B**), including neural cells of different stages of maturation, such as NPCs and newborn neurons (**Fig. 5C-E**). Clear differences in cell type composition were observed between the day 15 and day 30 organoids. Day 30 organoids contained more neurons and fewer NPCs, which further demonstrates the presence of more mature cell types in these organoids. We found no apparent difference in the contribution to the different clusters by *LINC00662*-CRISPRi organoids, suggesting that LINC00662 does not influence developmental fate in unguided neural organoids (**Fig. 5A-B**).

**Figure 5.**
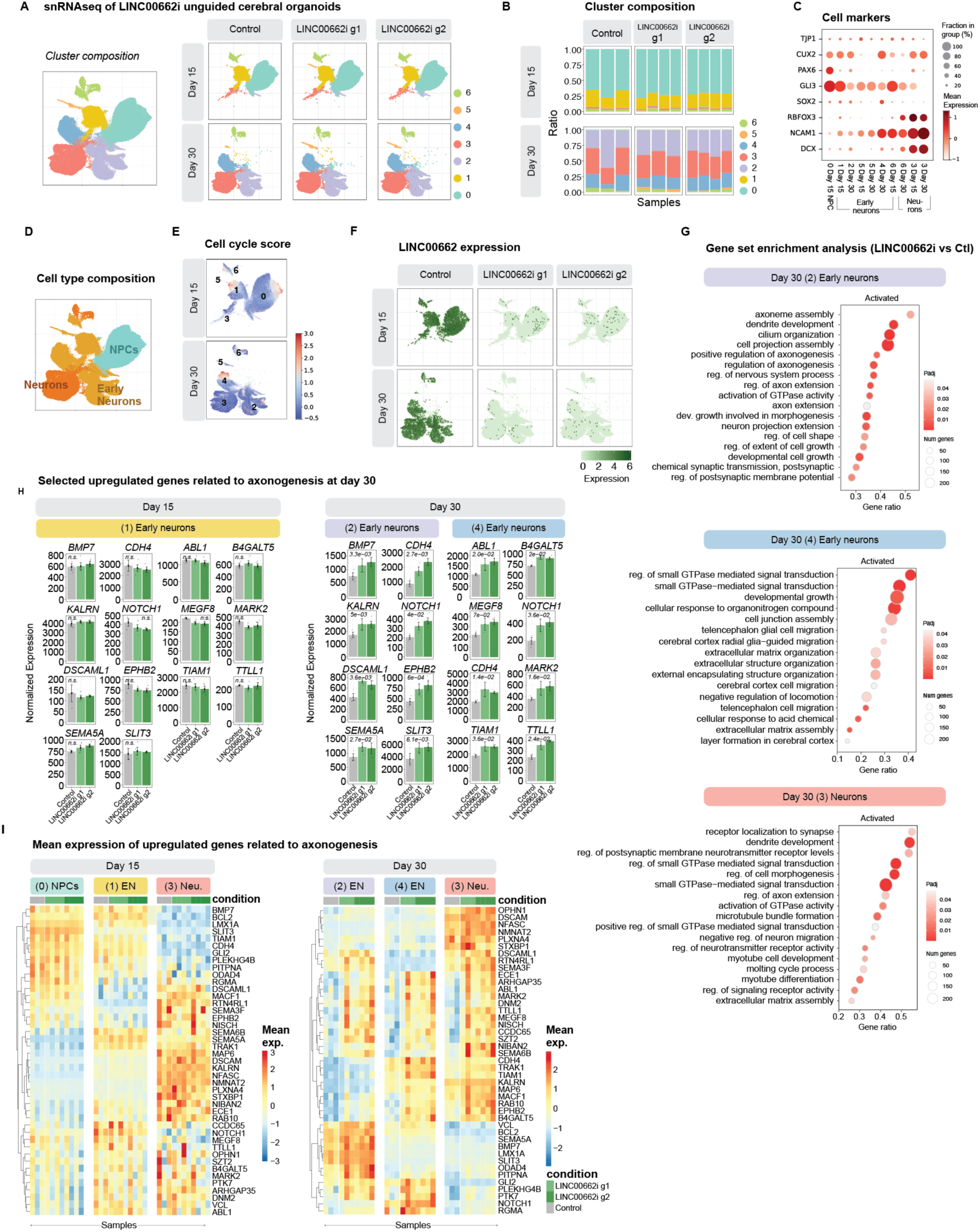
**A** UMAP embeddings of nuclei from day 15 and day 30 unguided neural organoids, showing clustering using all samples (left) and split per condition (right). **B** Barplot showing cluster composition per sample. **C** Expression of selected marker genes at both timepoints (day 15, day 30). Color indicates average expression per cluster; dot size indicates the percentage of nuclei in which the gene was detected. **D** UMAP coloured by annotated cell types. **E** UMAP showing cell cycle scores, calculated as the sum of gene module scores for S-phase and G2/M-phase markers (Scanpy:: score_genes_cell_cycle) **F** Expression of LINC00662 in day 15 (top) and day 30 (bottom) of control (left) and LINC00662-CRISPRi (right) unguided neural organoids according to snRNA-seq. **G** Selected enriched Gene Ontology (GO) terms from gene set enrichment analysis (GSEA) in early neurons (left and middle) and neurons (right) (clusterProfiler::gseGO). **H** Bar plots showing selected upregulated genes associated with enriched terms (from K) (Wald test, DESeq2). Each dot represents the summed expression in the corresponding samples’ cluster (cluster annotated at the top); bars show group mean with error bars representing the standard error of the mean. **I** Significantly upregulated genes among axon-related GO terms (from J) in early neurons or neurons (padj < 0.05, Wald test, DESeq2). Color indicates mean expression in each sample cluster (columns).

Next, we analyzed the transcriptional difference between control and *LINC00662*-CRISPRi organoids. We confirmed the transcriptional silencing of LINC00662 in all cell populations at both time points (**Fig. 5F**). To avoid pseudoreplication in our statistics, all downstream analyses were performed on pseudobulks of all nuclei in each sample cluster (summed counts). On day 15, we found very limited transcriptional differences between the *LINC00662*-CRISPRi and control organoids. This suggests that LINC00662 does not affect the earliest phase of neural differentiation, which is in line with the neurite extension analysis. However, in day 30 *LINC00662*-CRISPRi organoids we found changes in gene expression of many genes related to neuronal maturation and axon and dendrite formation. Gene Set Enrichment Analyses (GSEA) of the transcriptional differences identified in *LINC00662*-CRISPRi organoids compared to controls revealed a higher expression of genes related to terms linked to axonogenesis, axon extension, dendrite development, and cellular morphogenesis in three neuronal clusters of day 30 *LINC00662*-CRISPRi organoids (**Fig. 5G-I**). In line with this, we found that some of the upregulated genes in newborn neurons are key players in processes such as neurogenesis, e.g., NOTCH1 (Notch Receptor-1)^59^, neuronal connectivity, e.g., CDH4 (Cadherin-4)^60^, axon, and dendrite growth, e.g., SLIT3 (Slit Guidance Ligand 3)^61^ and KALRN (Kalirin RhoGEF Kinase)^62,63^ (**Fig. 5H-I**). The observed gene expression changes were very similar for the two different gRNAs. Together, these results demonstrate that CRISPRi-mediated silencing of LINC00662 leads to transcriptional changes in neural organoids on day 30 but not on day 15 of differentiation. These changes are characterized by increased expression of genes related to axonogenesis and dendrite formation. This confirms that the reduced neurite extension phenotype is due to a post-transcriptional mechanism and suggests that the increased expression of genes linked to this pathway is the consequence of a downstream compensatory mechanism. In summary, these findings further support the role of this lncRNA in controlling human neuronal maturation and neurite extension.

## Discussion

Retroviruses are infectious agents that can alter their genetic material and integrate into the host genome as proviruses resulting in the production of new virus copies. In some cases, retroviruses infect germline cells, allowing the integrated proviruses to be passed on to offspring as endogenous retroviruses^18,32^. PTERV1 likely entered the chimpanzee genome prior to the separation of humans, chimpanzees and gorillas, but the human genome was spared the subsequent expansion that occurred in chimpanzees and gorillas between 1.5 and 5 million years ago.^9^ This expansion of PTERV1 in the chimpanzee genome is likely to have had a significant impact on the population at that time.^9,35,36^. Endogenization of retroviruses is a relatively infrequent event over evolutionary timescales, making it challenging to observe directly and leaving many aspects poorly understood^64^. The ongoing endogenization of the koala retrovirus-A (KoRV-A) provides a contemporary parallel to historical events such as the PTERV1 invasion. KoRV-A is currently spreading among wild koala populations via horizontal transmission while simultaneously integrating into germline cells, thereby becoming a component of the inherited genome^65,66^. This process has had a major detrimental impact on the current koala population increasing disease burden, including a high prevalence of cancers^66^.

Our results support a scenario where PTERV1 infection and endogenization had two effects that influenced hominid speciation. First, it likely drove the ancestor of chimpanzees and bonobos through a population bottleneck^36^. Second, it resulted in more than 150 fixed PTERV1 insertions in the chimpanzee genome^9^ (**Fig. 1D**), some with the potential to affect gene regulatory networks, as exemplified here by *LINC00662*. Thus, our results provide new insights into how the colonization of primate genomes by retroviruses influences speciation, including the rewiring of gene regulatory networks during brain development.

Although millions of years have passed since most ERVs were integrated into the primate germline^67,68^, many ERVs contain sequences that can act as *cis*-acting regulatory elements^21,22^ As a result, and to protect against their mutagenic potential, genomes have evolved various defense mechanisms against TE invasion^69,70^. KRAB-ZNFs (KZNFs) are the largest family of transcription factors in primate genomes, and their rapid expansion in mammalian genomes correlates with the expansion of TEs^71,72^. The majority of KZNFs are DNA-binding transcriptional repressor proteins that bind to TEs. Their conserved N-terminal KRAB domain interacts with the epigenetic co-repressor TRIM28, which induces heterochromatin formation, ultimately leading to the accumulation of DNA methylation that represses the transcriptional and regulatory potential of ERVs and other TEs^23,24,26,71,73,74^. However, a consequence of this mechanism is the formation of new local mini-heterochromatin domains across species-specific TEs. In this study, we demonstrate that all 158 PTERV1 insertions in the chimpanzee genome are covered by DNA methylation in iPSCs and neural organoids. While the KZNF responsible for targeting PTERV1 has not yet been identified, our results support a model in which PTERV1s are recognized by the KZNF/TRIM28 repression machinery during early development, resulting in the establishment of DNA methylation across these loci. Importantly, these local heterochromatin domains are likely to influence gene expression in the vicinity, creating species-specific transcriptional patterns. We identified several transcripts that overlap with a PTERV1-insertion and exhibit differential expression in human and chimpanzee organoids. These include non-coding transcripts such as LINC00662, as well as protein coding genes such as for example, the GABA A receptor subunit beta-1 (GABRB1), which participates in synaptic transmission and is implicated in the pathogenesis of schizophrenia,^75,76^ and the low-density lipoprotein receptor-related protein 1b (LRP1B), which has been linked to several malignancies^77^. Thus, the impact of PTERV1s on the gene regulatory network in chimpanzee cells may be broad and not only limited to *LINC00662.* In support of this idea, the establishment of heterochromatin domains and spread of DNA methylation into adjacent genomic regions have also been documented for other classes of repetitive elements across diverse species^6,25,78–81^.

LINC00662 is a lncRNA that is highly expressed during human brain development. We found that in human NPCs, the promoter region of *LINC00662* is hypomethylated and decorated with the active histone mark H3K4me3, consistent with transcriptional activity. In contrast, chimpanzee NPCs exhibit no evidence of *LINC00662* transcription as indicated by the absence of H3K4me3 at the promoter and instead the presence of DNA methylation. This methylated state is likely a consequence of the PTERV1 insertion within intron 1 of *LINC00662* which may facilitate the spread of DNA methylation into the promoter region of *LINC00662*. Notably, LINC00662 expression in chimpanzee cells can be rescued by CRISPR deletion of the PTERV1 element, confirming that the presence of the PTERV1 prevents transcription. In summary, our data suggest that the PTERV1 insertion exerts a repressive influence on the *LINC00662* promoter, but further molecular studies are required to elucidate the precise mechanisms by which the PTERV1 insertion mediates this regulatory effect.

Our data demonstrate that LINC00662 is a human-specific noncoding transcript with high expression levels during brain development. While it is challenging to fully characterize the role of a human-specific lncRNA using *in vitro* organoid modelling, our results support a role for LINC00662 in the cytoplasm, where it binds to several cytoskeletal proteins involved in neural maturation and axon growth in the developing brain. Consistent with this, our loss-of-function experiments confirms that LINC00662 mediates a post-transcriptional mechanism that regulates neurite extension during the early differentiation of committed neuroblasts. Notably, our snRNA-seq analysis demonstrates that many transcripts related to these processes are upregulated in immature neurons in *LINC00662*-CRISPRi neural organoids, indicating a downstream feedback loop that activates these transcriptional programs. We do not fully understand the mechanisms underlying this observation, but it could be due a compensatory mechanism resulting from impaired neurite extension in *LINC00662*-CRISPRi organoids. Our findings provide evidence that post-transcriptional mechanisms may play an important role in the evolution of the human brain. Such mechanisms are challenging to study using transcriptome-based analyses, which currently limits the interpretation of many comparative experiments performed using iPSC-derived neurons or organoids from humans, chimpanzees, and other primate species. Nevertheless, our results provide further support for the idea that differences in dendrite and synapse maturation play a key role in mediating human-specific brain functions ^82,83^.

In summary, we demonstrate here how a chimpanzee-specific endogenous retroviral insertion contributes to transcriptional divergence in the development of the human and chimpanzee brains. Our results support the idea that ERVs and other TEs played an important role in the evolution of the hominoid brain.

## Methods

### Identification of full-length PTERV1s and genomic location analysis

The RetroTector software^38^ (https://github.com/PatricJernLab/RetroTector) was used to mine the chimpanzee genome (version Clint_PTRv2/panTro6 downloaded from https://hgdownload.soe.ucsc.edu/) for ERVs as previously described^84^. Predicted positions and structures for ERVs scoring 300 and above are summarized in Table S1.

To assess the genomic location (intragenic or intergenic) and orientation (sense or antisense) of PTERV1 elements relative to genes, transcript-level annotation was utilized. RetroTector output was intersected with the panTro6 RepeatMasker annotation (panTro6.fa.out, downloaded from https://hgdownload.soe.ucsc.edu/) using BEDTools (v2.26.0), to identify PTERV1 elements. Strand information was calculated based on the start and end coordinates of the PTERV1. For transcript-level annotation, the panTro6.ncbiRefSeq.gtf file (downloaded from https://hgdownload.soe.ucsc.edu/) was used. Features annotated as "transcript" were filtered, strand information was retained, and only annotations corresponding to canonical chromosomes 1, 2A, 2B, 3–22, X, and Y were included in the analysis.

To identify intragenic PTERV1 elements, BEDTools (v2.26.0) was employed using the intersect command with the parameters -a, -b, -wa, and -wb. The resulting overlaps were processed using an awk script to compare the strand orientations of the intersecting features. A new column was added to the output to indicate whether the PTERV1 element and the associated gene were in the "same" or "opposite" orientation. Finally, strand orientation frequencies were summarized using an additional awk script to count the occurrences of "same" and "opposite" orientations, avoiding double-counting overlapping PTERV1s.

To identify homologous genes containing PTERV1 insertions in chimpanzee, BEDTools intersect (v2.30.0) was used to compare PanTro6 RefSeq gene annotations (-a) and RetroTector-predicted PTERV1 annotation (-b). Coordinates of genes hosting an entire PTERV1 insertion (-wa -F 1) without an annotated homologue in the human reference genome version hg38, were lifted to hg38 coordinates using UCSC LiftOver web browser tool.

Finally, the resulting lifted coordinates (-a) were intersected with human GENCODE v38 gene annotation (-b) using BEDTools intersect with default parameters (-wo). This analysis identified 26 genes annotated in hg38 with homologous to pantro6 genes containing PTERV1 insertions. These candidates were further scrutinized based on their difference of expression between human and chimpanzee organoids using bulk RNA-seq (see Methods: *Bulk RNA-seq analysis of human and chimpanzee organoid samples*).

### iPSC culture

The two chimpanzee iPSC lines used in this study have been generated in other labs and previously reported: Sandra A (female)^41^, generated with non-integrative episomal vectors, here referred to as PT1, Jo_C (male)^40^, generated with non-integrative episomal vectors, here referred to as PT2. The two human induced pluripotent stem cell (iPSC) lines used in this study were obtained from RIKEN. These lines were generated using non-integrative episomal vectors^85^: RBRC-HPS0328 606A1 (female), here referred to as HS1 (RIKEN, RRID:CVCL_DQ11), RBRC-HPS0331 610B1 (male), here referred to as HS2 (RIKEN, RRID: CVCL_DQ4).

The iPSC lines were maintained on LN521 (0.7 *µ*g/cm2; Biolamina) coated Nunc multidishes in iPS media (StemMACS iPS-Brew XF and 0.5% penicillin/streptomycin; GIBCO). iPSCs were passaged 1:2-1:6 every 2-3 days when 70-90% confluency was reached. Media change was performed daily and 10 *μ*M Y27632 (Rock inhibitor, Miltenyi) was added when cells were passaged.

### Unguided neural organoid culture

Human and chimpanzee unguided neural organoids were generated using the STEMdiff^TM^ Cerebral Organoid kit (STEMCELL Technologies) following the manufacturer’s instructions with some adjustments. Briefly, human (HS1 and HS2) and chimpanzee (PT1 and PT2) iPSCs were maintained in StemMACS iPS brew XF as described under “iPSC culture”. Chimpanzee iPSCs were transferred to mTeSR1 medium (STEMCELL Technologies) two weeks before starting the organoids and maintained as described under “iPSC culture” with mTeSR1 replacing StemMACS iPS brew XF. After that, the timing of the STEMdiff^TM^ Cerebral Organoid kit protocol was followed with the exception of PT2, which was transferred to neural induction medium one day earlier than described. In addition, EBs for HS1, HS2 and PT1 were generated using the kit’s EB seeding medium, but EBs for PT2 using mTeSR1. Briefly, at day 0 of differentiation, iPSCs at 70-90% confluency were detached and centrifuged, then resuspended in 1 ml of EB Seeding Medium (STEMdiff^TM^ Cerebral Organoid Basal Medium 1 and STEMdiff^TM^ Cerebral Organoid Supplement A, 4:1, supplemented with 10 µM of Y-27632 (Rock Inhibitor)) or mTeSR1 (supplemented with 10 µM Y-27632 (Rock Inhibitor)). 9000 cells per well in 100 µl EB seeding medium (HS1, HS2, PT1) or 10000 cells in 250 µl mTeSR1 (supplemented with 10 µM Y-27632 (Rock Inhibitor); PT2) were then distributed in a round-bottom ultra-low attachment 96-well plate. EBs were incubated at 37°C for 48 h. At day 2 and 4 of differentiation 100 µl of fresh EB Formation Medium (STEMdiff^TM^ Cerebral Organoid Basal Medium 1 and STEMdiff^TM^ Cerebral Organoid Supplement A, 4:1; HS1, HS2, PT1) or 200 µl of mTeSR1 (PT2) were added to each well. On day 5, or when the organoids reached a diameter of >300 µm, organoids were transferred to an ultra-low attachment 24-well plate with 0.5 ml of Induction Medium (STEMdiff^TM^ Cerebral Organoid Basal Medium 1 and STEMdiff^TM^ Cerebral Organoid Supplement B) in each well (1 organoid/well to avoid organoid fusion). Organoids were incubated at 37°C for 48 h. On day 7, organoids were embedded with ∼15 µl of Matrigel per organoid on an embedding surface (Parafilm) in a sterile 100 mm dish and incubated at 37°C for 30 min to polymerize the Matrigel. Organoids were then transferred to an ultra-low attachment 6-well plate with 3 ml of Expansion Medium (STEMdiff^TM^ Cerebral Organoid Basal Medium 2, STEMdiff^TM^ Cerebral Organoid Supplement C, STEMdiff^TM^ Cerebral Organoid Supplement D) per well. 10-12 organoids per well were cultured together. On day 10, the organoids were transferred to an orbital shaker (rpm = (√RCF/throw * 1.118) * 1000), and media was changed with 3 ml/well of Maturation Medium (STEMdiff^TM^ Cerebral Organoid Basal Medium 2, STEMdiff^TM^ Cerebral Organoid Supplement E). The medium was replaced every 3 days. Two independent batches of organoids per cell line were generated. On day 15 of differentiation, the organoids were collected for downstream analyses. 5 organoids were collected and snap-frozen for each bulk RNA sequencing replicate (n = 3 replicates per cell line per batch). 5 organoids were collected and snap-frozen for each snRNA-seq replicate (n = 1 replicate per cell line per batch). 3 organoids were collected for immunostaining (see: Immunohistochemistry). 5 organoids were collected for long-read DNA sequencing.

### Immunocytochemistry

The iPSCs were washed two times with KPBS for 5 min and subsequently fixed with 4% paraformaldehyde for 15 min at room temperature. This was followed by two washes with KPBS for 5 min. The cells were blocked and permeabilized in blocking solution (0.3% Triton X-100 and 5% normal donkey serum in KPBS) for 45 min at room temperature. The primary antibody (rabbit anti-NANOG (Abcam AB446437), 1:1000) was added in blocking solution and incubated overnight. On the next day, the cells were washed two times with KPBS for 5 min. The secondary antibody (donkey anti-rabbit Alexa Fluor 488 (Invitrogen A21206), 1:500; or donkey anti-rabbit Cy5 (Jackson ImmunoResearch AB_2340607), 1:500) was added in blocking solution and incubated at room temperature for 1 hour followed by three washes with KPBS for 5 min. DAPI (1:1000; Sigma-Aldrich) in KPBS was added as a nuclear counterstain and incubated for 5 min at room temperature. The cells were washed two times with KPBS for 5 min and imaged on a Fluorescence Microscope Leica DMI6000B.

### Immunohistochemistry

Unguided neural organoids were fixed in 4% paraformaldehyde for 1 hour at room temperature. After that, they were washed three times with KPBS and left in a 1:1 30% sucrose solution and OCT (catalog no. 45830, HistoLab) mixture overnight at 4°C. Organoids were then transferred to a cryomold containing OCT, frozen on dry ice, and stored at −80°C in freezer bags.

Ahead of staining, organoids were sectioned on a cryostat at −20°C at a thickness of 20 µm and placed onto Superfrost plus microscope slides. They were then washed two times with KPBS for 5 min and subsequently blocked and permeabilized in 0.25% Triton X-100 and 4% normal donkey serum in KPBS for 1 hour at room temperature. The primary antibodies [rabbit anti-PAX6 (BioLegend, 901301), 1:500 dilution; mouse anti-ZO1 (Invitrogen, 339100), 1:300 dilution; rabbit anti-beta tubulin III (Biolegend, 802001), 1:1000 dilution; and goat anti-SOX2 (R&D Systems, AF2018)] were prepared in antibody solution (0.1% Triton X-100 and 4% normal donkey serum in KPBS), added to the sections and incubated overnight at 4°C. Subsequently, the sections were washed two times with KPBS and one time with wash solution (0.5% Triton X-100 in KPBS). The secondary antibody [donkey anti-rabbit Alexa Fluor 488 (Invitrogen A21206), 1:1000; or donkey anti-rabbit Cy3 (Jackson ImmunoResearch AB_2307443), 1:400; donkey anti-goat Alexa Fluor 488 (Invitrogen A11055), 1:500; donkey anti-rabbit Alexa Fluor 568 (Invitrogen A10042), 1:500; and donkey anti-mouse Alexa Fluor 647 (Invitrogen A31571), 1:1000] was prepared in antibody solution, added to the sections and incubated at room temperature for 1 hour protected from light, followed by two washes with KPBS and one wash with washing solution. The sections were incubated with DAPI in KPBS (1:1000) for 5 min at room temperature and washed once with KPBS. The sections were mounted using FluorSave (Merck, 345789) and imaged using an Operetta CLS (Perkin Elmer) or on a Fluorescence Microscope Leica DMI6000B. Figures were assembled in ImageJ (NIH) and Adobe Photoshop (Adobe Systems, San Jose, CA); only contrast and brightness were adjusted to optimize the image quality.

Mean fluorescence intensities of DAPI and TUJ1 were measured for 2-3 regions of interest (rosette and intermediate zone) from six organoids derived from two independent batches at 15 and 30 days. Regions were chosen manually in a blinded manner based on DAPI staining after 1) manually marking the rosette ventricle followed by automatic expansion of the region with 18pixels leaving a donut shaped ROI and 2) manually drawing intermediate zone following the rosette edge using ImageJ (NIH). To control for variation in cell number across sections, TUJ1 values were normalized to the corresponding mean DAPI fluorescence intensity for each ROI. Two-tailed Student’s *t* test, or one-way ANOVA followed by Dunnett’s test using α = 0.5 was conducted. Statistical analysis was performed in GraphPad Prism (GraphPad Software, San Diego, USA).

### Single-nuclei RNA-seq

Nuclei isolation from organoids was performed as described previously^86^. Briefly, the organoids were thawed and dissociated in ice-cold lysis buffer [0.32 M sucrose, 5 mM CaCl_2_, 3 mM MgAc, 0.1 mM Na_2_EDTA, 10 mM Tris-HCl (pH 8.0), and 1 mM dithiothreitol] using a 1-ml tissue douncer (Wheaton). The homogenate was carefully layered on top of a sucrose solution [1.8 M sucrose, 3 mM MgAc, 10 mM Tris-HCl (pH 8.0), and 1 mM dithiothreitol] and centrifuged at 30000 × g for 2 hours and 15 min. Pelleted nuclei were softened for 10 min in 100 µl of nuclear storage buffer [15% sucrose, 10 mM Tris-HCl (pH 7.2), 70 mM KCl, and 2 mM MgCl_2_], resuspended in 300 µl of dilution buffer [10 mM Tris-HCl (pH 7.2), 70 mM KCl, and 2 mM MgCl_2_] and ran through a cell strainer (70 µm). The nuclei were sorted via FANS (with a FACS Aria, BD Biosciences) at 4° C at low flow rate using a 100 μm nozzle (reanalysis showed >95% purity).

FACS-sorted nuclei (8500 - 10000 nuclei/cells per sample) were directly loaded onto the Chromium Next GEM Chip G Single Cell Kit along with the reverse transcription mastermix following the manufacturer’s protocol for the Chromium Next GEM single cell 3′ kit (PN-1000268, 10X Genomics) to generate single-cell gel beads in emulsion. cDNA amplification was done according to the guidelines from 10X Genomics using 13 cycles of amplification for 3′ libraries. Sequencing libraries were generated with unique dual indices (TT set A) and pooled for sequencing on a Novaseq6000 or Novaseq X Plus using a 100-cycle kit and 28-10-10-90 reads.

#### snRNA-seq analysis

Basecalling of reads and generation of sample-specific FASTQ files were performed using 10X Genomics mkfastq (v6.0.0; RRID:SCR_017344). Alignment to the reference genome (hg38) and gene quantification was performed using 10X Genomics Cell Ranger count (v6.0.0) including intronic reads (--include-introns).

#### snRNA-seq analysis of human and chimpanzee organoids

##### Quality Control

Preprocessing and quality control of all samples were conducted using Seurat (v5.1.0). Read count matrices (produced by Cell Ranger) were loaded into individual Seurat objects using Seurat::Read10X. Genes expressed in fewer than 3 nuclei in the whole dataset were excluded when creating the individual Seurat objects (Seurat::CreateSeuratObject, min.cells = 3). Nuclei containing more than 2% mitochondrial gene counts, with less than 500 genes detected, or with a number of genes detected over two standard deviations, or under one standard deviation from the sample’s mean were excluded from further analyses.

##### Clustering and UMAP Embeddings

Samples’ objects were integrated using Seurat::FindIntegrationAnchors (dims = 1:30) and Seurat::IntegrateData (dims = 1:30). Count scaling was performed using Seurat::ScaleData across all genes. Principal component analysis (PCA) was conducted using Seurat::RunPCA using the most variable features on the dataset (Seurat::VariableFeatures). Neighborhood graph construction was based on 10 PCAs and 20 nearest neighbors (Seurat::FindNeighbors, dims = 1:10, k.param = 20). Clustering was performed using Seurat::FindClusters with a resolution of 0.1 (res). Uniform Manifold Approximation and Projection (UMAP) reduction was calculated using Seurat::RunUMAP (dims = 1:10).

##### Cell Type Annotation

Cell cycle scores were calculated using a previously established list of cell cycle-related genes using Seurat::CellCycleScoring. Cell type annotations were manually assigned based on the expression of canonical gene markers.

#### snRNA-seq analysis of human LINC00662-CRISPRi organoids

##### Quality Control

Preprocessing and quality control of all samples were conducted using Scanpy (v1.10.1). Read count matrices (produced by Cell Ranger) were loaded into individual AnnData objects using Scanpy::read_10x_mtx. Genes expressed in fewer than 3 nuclei in the whole dataset were excluded using Scanpy::filter_genes (min_cells=3). Quality control metrics were computed with Scanpy::calculate_qc_metrics, considering mitochondrial and ribosomal gene content (percent_top=None, log1p=False). Doublets identified by Scanpy::Scrublet were excluded from further analyses. Nuclei containing more than 2% mitochondrial gene counts, with less than 500 genes detected, or with a number of genes detected outside two standard deviations from the sample’s mean, were excluded from further analyses.

##### Clustering and UMAP Embeddings

Samples were merged using Anndata::concat (join=’outer’). Count normalization was performed with Scanpy::pp.normalize_total (target sum = 1e6), followed by natural log transformation (Scanpy::pp.log1p) and scaling (Scanpy::pp.scale, max_value=10).

Principal component analysis (PCA) was conducted using arpack solver (Scanpy::tl.pca, svd_solver=’arpack’). Neighborhood graph construction was based on 20 PCAs and 15 nearest neighbors (Scanpy::pp.neighbors, n_neighbors = 10, n_pcs = 20). Clustering was performed using Leiden algorithm (Scanpy::tl.leiden, resolution=0.1, directed=False, flavor=’igraph’).

A graph abstraction was generated via Partition-based Graph Abstraction (PAGA) (Scanpy::tl.paga), and the Uniform Manifold Approximation and Projection (UMAP) layout was computed using PAGA-initialized positions (Scanpy::tl.umap, init_pos=’paga’). Upon examination of UMAP embeddings, batch effects (e.g., experimental organoid batches) were integrated using Harmony (Scanpy::external.pp.harmony_integrate). The nearest-neighbor graph was then recomputed using the Harmony-adjusted PCA (Scanpy::pp.neighbors, use_rep=’X_pca_harmony’), followed by re-clustering with Leiden and re-computation of PAGA and UMAP using the same paramaters as above.

##### Cell Type Annotation

Cell cycle scores were calculated using a previously established list of cell cycle-related genes using Scanpy::tl.score_genes_cell_cycle. Cell type annotations were manually assigned based on the expression of canonical gene markers.

##### Pseudobulks and gene differential expression analyses

Raw gene counts were summed per cluster per sample, to produce seven (clusters 0 to 6 as shown in **Fig 4D**) cluster-specific gene count matrices where each column is a sample, and each gene is a row. These pseudobulked gene count matrices were input to DESeq2 with a pseudocount of 1, to perform differential expression analyses between LINC00662-CRISPRi and Control organoids per timepoint (day 15, day 30), per cluster (clusters 0-6). Design for the test was set to ∼ batch + condition, to test for transcriptional differences between conditions (LINC00662-CRISPRi / Control) while accounting for experimental organoid batches. Log2FoldChanges were shrinked using DESeq2::lfcShrink.

Gene log2FoldChanges were ranked and input as a named list to clusterProfiler::gseGO from which a minimum gene set size was set to 3 (minGSSize) and maximum of 800 (maxGSSize). Bonferroni and Hochberg correction (pAdjustMethod = “BH”) was performed on the results and a p-value cut-off of 0.05 (pvalueCutoff) was used to filter significant terms. Heatmap visualization (**Fig 4K**) was performed on mean normalized counts (DESeq2::counts, normalized = T) using pheatmap scaling per gene (scale = “row”).

### Bulk RNA-seq

Total RNA was isolated from organoids using the RNeasy Mini Kit (QIAGEN). Libraries were generated using Illumina TruSeq Stranded mRNA library prep kit [poly(A) selection] and sequenced on a Novaseq 6000 or Novaseq X Plus (PE, 2 × 150 bp).

#### Bulk RNA-seq analysis of human and chimpanzee organoid samples

Reads were aligned to the reference genome (hg38) using STAR aligner (v2.7.8a) using a unique mapping approach (--outFilterMultimapNmax 1) with a maximum allowed mismatch of 3% (--outFilterMismatchNoverLmax 0.03). Alignment files were sorted and indexed using SAMtools (v1.16.1) and input to deeptools BAMCoverage function (v2.5.4) splitting strands (--filterRNAstrand) and normalizing using RPKM (--normalizeUsing RPKM).

##### Gene differential expression analysis

Gene count matrices of Gencode v38 were produced using featureCounts (Subread package, v1.6.3, -s 2). Differential expression analysis between the two species was performed using DESeq2 (v1.44.0, design = ∼ species). Log2FoldChanges were shrinked using DESeq2::lfcShrink. Heatmap visualization (**Fig 2B**) was performed on mean normalized counts (DESeq2::counts, normalized = T) using pheatmap (scale = “none”).

### Long-read DNA sequencing

High molecular weight (HMW) DNA was extracted from frozen human fetal brain tissue, human and chimpanzee iPSC pellets (500000 - 1 million cells) and day 15 unguided neural organoids (5 organoids) using the Nanobind HMW DNA Extraction kit (PacBio) following the manufacturer’s instructions. HMW DNA was eluted in 100 µL of Buffer EB. Size selection to enrich for fragments >5 kb was performed using the Short Read Eliminator XS kit (PacBio) according to the manufacturer’s instructions. HMW DNA quantity and quality were assessed by Nanodrop and Qubit from the top, middle, and bottom of each tube, and on a TapeStation 4200 (Agilent) using the Agilent Genomic DNA Screen Tape Assay (Ref No. G2991-90040, Edition 08/2015). HMW DNA was sheared to approximately 10kb fragment length using a g-TUBE (Covaris) following the manufacturer’s protocol. Library preparation for whole genome sequencing was performed using the SQK-LSK114 Ligation Sequencing kit (Oxford Nanopore Technologies) with >1 mg DNA input. Libraries were each sequenced separately using a FLO-PRO114M PromethION Flow Cell R10.4.1 on a PromethION platform (Oxford Nanopore Technologies) at SciLifeLab Uppsala or in-house on a PromethION 2 Solo (Oxford Nanopore Technologies) for 72 hours.

#### Long-read DNA analysis

Raw sequencing data (POD5 format) were basecalled and mapped to either the human reference genome (GRCh38/hg38) or chimpanzee reference genome (Clint_PTRv2/panTro6) using Dorado version 0.5.1-CUDA-11.7.0 (https://github.com/nanoporetech/dorado) utilizing the super accurate basecalling model <u>dna_r10.4.1_e8.2_400bps_sup@v4.3.0</u> and <u>dna_r10.4.1_e8.2_400bps_sup@v4.3.0_5mCG_5hmCG@v1</u> for methylation aware basecalling. Resulting BAM files were sorted and indexed using SAMtools version 1.18^87^. PTERV1 and LINC00662 methylation were visualized with MethylArtist version 1.2.4^88^ using the segmeth (parameters:–excl_ambig), segplot and locus functions with default parameters.

### Long-read direct RNA sequencing

#### RNA extraction

RNA was extracted from fetal forebrain samples using the TRIzol RNA extraction protocol (Cat #15596026). Briefly, the tissue was homogenized in 1 ml of TRIzol Reagent (Invitrogen; Pub. No. MAN0001271). 200 μl of chloroform were then added and mixed to the lysate after 5 minutes of incubation at room temperature (RT). After 3 minutes of incubation (RT), the sample was centrifugated at 12000 × g at 4 °C for 20 minutes. The aqueous phase (containing the RNA) was transferred to a new tube and 500 μl of isopropanol were added followed by a 10 minutes incubation at 4 °C. RNA was then pelleted by centrifugation at 12000 × g for 10 minutes at 4 °C. RNA was resuspended in 1 ml of 75% ethanol, vortexed, and centrifuged for 5 minutes at 7500 × g at 4 °C. The RNA pellet was air-dried and resuspended in 35 μl of RNase-free water. The suspension was then incubated at 56 °C for 15 minutes. DNase treatment (Pub. No 1907M) was then performed on the isolated RNA using the TURBO DNA-*free*^TM^ Kit (Invitrogen, Cat #AM1907) according to manufacturer’s instructions. Poly(A) mRNA isolation, library preparation and sequencing were performed as previously described^89^.

#### Long-read direct RNA analysis

Basecalling was performed using dorado (version 0.7.1-CUDA-11.7.0) with model rna004_130bps_sup@v3.0.1 and modified bases model rna004_130bps_sup@v3.0.1_m6A_GRACH@v1 (--modified-bases-models). Reads were mapped to the reference genome (hg38) using minimap2 (-ax splice -uf -k14 -y). Genome browser visualization was done in IgV (version 2.18.2) (http://www.broadinstitute.org/igv/) using the sorted and indexed alignment files (samtools 1.16.1).

### Generation of *LINC00662-*PTERV1 Knockout chimpanzee iPSCs

#### Lentiviral production

Lentivirus was produced as previously described^90^. HEK293T cells (RRID:CVCL_0063) at a confluency of 70-90% were transfected with third-generation packaging and envelope vectors pMDL (Addgene #12251), psRev (Addgene #12253), and pMD2G (Addgene #12259) with Polyethyleneimine (PEI Polysciences PN 23966) in DPBS (GIBCO). Lentivirus was harvested 48h after transfection by filtering and centrifugation at 25000 × g for 1.5h at 4°C. Virus was resuspended in DPBS, aliquoted and stored in −80°C until further use. Lentivirus were in titers of 10^9^, determined by standard qRT-PCR.

#### CRISPR-Cas9 gene editing

Chimpanzee cell line PT1 was dissociated using Accutase and 200000 cells were plated in laminin-521 coated wells in iPS media with 10 μM RY27632. Cells were transduced with lentiviral vectors pLV.U6.EFS-NS.H2B-RFPW (Addgene plasmid # 170363; RRID:Addgene_170363) with MOI 1 and pLV.CAS9-GFP (Addgene plasmid # 170361; RRID:Addgene_170361) with MOI 20 carrying sgRNA1 and sgRNA2 respectively, designed to target the flanking regions of PTERV1. Cells were expanded for 10 days prior to FACS (FACSAriaIII, BD sciences). Cells were detached using Accutase and resuspended in iPS media containing RY27632 (10 μM) and DRAQ7 (1:1000). Gating parameters were determined by side and forward scatter to eliminate debris and aggregated cells. The GFP^+^ and RFP^+^ gates were set using untransduced and single transduced iPSCs. The sorting gates and strategies were validated via reanalysis of sorted cells (> 99% purity cut-off). 300000 RFP^+^/GFP^+^/DRAQ7^-^ cells were collected, spun down at 400*g* for 5 minutes and kept at −80°C until further analysis.

sgRNA1: GCAGCACTGCGCATCTCTGA
sgRNA2: ACCAGGTAATGAAAAGAGCC

To confirm successful excision of PtERV1, genomic DNA was extracted from WT and PTERV1-excised PT1 cells according to manufacturer’s instructions (DNeasy Blood and Tissue kit, Qiagen). Using genomic primers targeting LINC00662, a 7500 bp region was amplified (PrimeSTAR GXL DNA Polymerase, TAKARA). The PCR product of the expected size was purified (QIAquick Gel Extraction Kit, Qiagen) and confirmed through automated Sanger sequencing (EuroFins genomics).

gPCR_LINC00662_Fwd: GCGGCTGATCTCACCTTGTA
gPCR_LINC00662_Rev: ACGCAGCAGGACAGAATCTC

### qRT-PCR

Total RNA isolated from iPSCs or unguided neural organoids (RNeasy mini kit, Qiagen) was reverse transcribed using the Maxima cDNA synthesis kit (Thermo Scientific). qRT-PCR was performed using primers listed below and SYBR green master mix (Roche) on a LightCycler® 480. Three technical replicates per sample were amplified for 40 cycles and cycle values were normalized to ß-actin and HPRT.

Primer sequence (5′to 3′)
LINC00662 Fwd: CAGGCCACTTACAGACTCCA
LINC00662 Rev: AGTGTTTCCAGCCTGAGACT
ß-actin Fwd: CCTTGCACATGCCGGAG
ß-actin Rev: GCACAGAGCCTCGCCTT
HPRT Fwd: ACCCTTTCCAAATCCTCAGC
HPRT Rev: GTTATGGCGACCCGCAG

### fbNPC differentiation

iPSCs differentiation into forebrain neural progenitors (fbNPCs) was performed as previously described^6,91^. Briefly, iPSCs at 70-90% confluency were dissociated using Accutase and plated in N2 differentiation medium [1:1 DMEM/F-12 (21331020; GIBCO) and Neurobasal (21103049; GIBCO) supplemented with 1% N2 (GIBCO), 2 mM L-glutamine (GIBCO), and 0.2% penicillin/streptomycin] on LN111-coated (1.14µg/cm^2^; Biolamina) Nunc Δ multidishes at a density of 10000 cells/cm^2^ with 10 μM SB431542 (Axon) and 100 ng/ml noggin (Miltenyi) for dual SMAD inhibition, and 10 μM Y27632. Media (N2 medium + SB + noggin) was changed every 2-3 days. On day 9 of differentiation, media was changed to N2 media without SMAD inhibitors. On day 11, cells were replated in B27 medium [Neurobasal supplemented with 1% B27 without vitamin A (GIBCO), 2 mM L-glutamine, 0.2% penicillin/streptomycin, BDNF (20 ng/ml; R&D), L-ascorbic acid (0.2 mM; Sigma), and Y27632 (10 μM)] on LN111-coated Nunc Δ multidishes at a density of 800000 cells/cm^2^. Cells were harvested at day 14 for downstream analysis.

### Cellular fractionation assay

Cell fractionation was performed using the Paris^TM^ kit (Thermo Fisher; Part Number AM1921) on HS1 iPSC-derived fbNPCs. Briefly, nuclear and cytoplasmic fractions were separated from fresh cultured cells. In parallel, total RNA was extracted from whole cell lysate from the same culture. RNA was isolated from whole cells, nuclear and cytoplasmic lysate individually according to manufacturer’s instructions. Isolated RNA was reversed transcribed into cDNA and analyzed using qRT-PCR as described above.

### Chop mass spectrometry

This experimental workflow was based on the ChIRP-MS protocol published earlier^92^. HS1 iPSCs were used to generate fbNPCs as described above. On day 14, fbNPCs were trypsinised, washed with DPBS and spun down at 500 × g for 5 minutes. Cells were resuspended in 1% formaldehyde in DPBS and kept at 1-2.5×10^6^cell/mL of 1% formaldehyde. Cells were incubated at room temperature for 30 minutes with end-to-end rotation. The reaction was stopped with 0.125mM glycine for 5 minutes with end-to-end rotation. Cells were pelleted at 4000 × g for 5 minutes before being rinsed with DPBS and pelleted again. The supernatant was discarded, and the pellet was processed for DNase digestion.

To digest genomic DNA, pellets were treated with 200 µL of TURBO nuclear lysis buffer (50 mM Tris pH 7.0, 0.5% NP-40, 1× Turbo DNase buffer, 1× protease inhibitor cocktail, 1 mM PMSF, 10 U/mL TURBO DNase, and 0.1 U/μL RNase inhibitor), followed by the addition of 10 µL of enzyme per sample. Incubation was carried out at 37°C for 30 min. Reactions were stopped by adding 850 µL of STOP lysis buffer (50 mM Tris pH 7.0, 50 mM EDTA, 50 mM EGTA, 1% SDS, 0.5% NP-40, 1 mM DTT, 1 mM PMSF, protease inhibitor, and 0.1 U/μL RNase inhibitor), followed by 15 min incubation on ice with intermittent pipetting. Nuclear homogenization was achieved by passing the lysate through a syringe needle.

Sonication was performed using Bioruptor with 30 ON/OFF cycles (or until clear solution). The chromatin was clarified by centrifugation (16000 × g, 10 min, 4°C), and the supernatant was pooled across replicates. Hybridization buffer (50 mM Tris pH 7.0, 10 mM EDTA, 1% SDS, 1 mM DTT, 750 mM NaCl, 15% formamide, 1 mM PMSF, protease inhibitors, and 0.1 U/μL RNase inhibitor) was added in a 2:1 ratio to the chromatin volume.

For pre-clearing, 100 µL of C1 beads per 100 million cells were washed twice with 500 µL nuclear lysis wash buffer (50 mM Tris pH 7.0, 10 mM EDTA, 1% SDS, 1 mM DTT, 1 mM PMSF, protease inhibitors, 0.1 U/μL RNase inhibitor) and incubated with the chromatin at 37°C for 30 min. The cleared supernatant was separated magnetically and distributed into individual tubes for pull-down reactions, reserving 20 µL for RNA, DNA, and protein inputs. Biotinylated oligonucleotide probes (1–2 µL of 100 µM pool per reaction) specific to LIN00662 and a LacZ control were added, followed by a 4 h hybridization at 37°C.

In parallel, streptavidin beads (110–210 µL) were washed twice in nuclear lysis wash buffer, blocked with tRNA (500 ng/µL) and BSA (1 mg/mL) for 1 h at 37°C, and washed once again. Post-hybridization, beads were added to the chromatin-probe mix and incubated for an additional hour at 37°C. The complexes were washed three times with wash buffer 1 (2× SSC, 0.5% SDS, 1 mM PMSF, 1 mM DTT, 0.1 U/μL RNase inhibitor) at 37°C, 500 rpm for 5 min each. After magnetic separation, residual buffer was removed by brief centrifugation.

For protein recovery, 100 µL of biotin elution buffer (7.5 mM HEPES, 12.5 mM biotin, 0.15% SDS, 0.075% sarkosyl, 0.02% sodium deoxycholate, 75 mM NaCl, 1.5 mM EDTA) was added and incubated for 20 min at room temperature with agitation, followed by 10 min at 65°C. The eluate was collected magnetically, and the process was repeated once for a final volume of 200 µL, reserving 4 µL (2%) for RNA RT-PCR analysis.

For RNA isolation, both input and bead-bound fractions were treated with elution buffer and proteinase K, incubated at 50°C for 50 min to reverse cross-links, and purified using Trizol followed by Zymo RNA Clean & Concentrator columns (Zymo Research, R1014). TURBO DNase digestion was performed to eliminate any residual genomic DNA, and enrichment of target transcripts was confirmed via RT-PCR.

#### Mass spectrometric analysis

Mass spectrometric analysis was carried out using an Orbitrap Fusion Tribrid instrument coupled to an Easy-nLC 1200 nano-liquid chromatography system (Thermo Fisher Scientific). Peptide samples were first loaded onto an Acclaim PepMap 100 C18 trap column (100 μm × 2 cm, 5 μm particle size; Thermo Fisher Scientific) and subsequently separated on an in-house packed analytical column (75 μm × 300 mm, 3 μm ReproSil-Pur C18, Dr. Maisch). A linear gradient from 5% to 28% acetonitrile (ACN) in 0.1% formic acid (FA) was applied over 75 minutes, followed by a rapid increase to 80% ACN in 0.1% FA for 5 minutes.

Full MS spectra were collected in the Orbitrap at a resolution of 120,000 within an m/z range of 375–1500. The most intense precursor ions (charge states 2–7) were selected for data-dependent fragmentation using higher-energy collisional dissociation (HCD) at a normalized collision energy of 30. The “top speed” duty cycle was maintained at 1 s, with an isolation window of 0.7 m/z and dynamic exclusion set to 10 ppm for 45 s. MS/MS scans were recorded at 30000 resolution with a maximum injection time of 110 ms. Analytical blanks were run between samples to prevent carryover.

Raw data were processed using Proteome Discoverer software (v2.4, Thermo Fisher Scientific). Peptide identification was performed with Mascot against the SwissProt human database, allowing one missed tryptic cleavage. Search parameters included precursor and fragment mass tolerances of 5 ppm and 30 mmu, respectively. Methionine oxidation and cysteine alkylation were set as variable modifications. False discovery rate (FDR) control was implemented using a decoy database search, applying a strict 1% threshold at the peptide-spectrum match (PSM) level. Identified proteins were grouped by shared peptide sequences to reduce redundancy and filtered at 5% FDR.

#### Chop mass spec analysis

All downstream analysis was done using R. Peptide counts in each replicate were subtracted by the corresponding values of LacZ control to remove background noise. Further, the data was normalized by dividing the corrected counts by the total count in the respective replicate and scaled to 1000. Based on average of normalized values across the replicates, the top 45 proteins were analyzed using StringDb (v11.5; RRID:SCR_005223)^93^. Additionally, information on the subcellular location of the proteins were obtained from the Human Protein Atlas (RRID:SCR_006710; proteinatlas.org)^94–96^.

### Generation of *LINC00662*-CRISPRi iPSCs and unguided neural organoids

To silence the expression of *LINC00662* in iPSCs, we used the protocol described in Johansson et al.^97^. Single guide sequences targeting near the transcription start site (TSS) were designed according to the GPP Portal (Broad Institute). The guide sequences were inserted into a deadCas9-KRAB-T2A-GFP lentiviral backbone, pLV hU6-sgRNA hUbC-dCas9-KRAB-T2a-GFP, a gift from Charles Gersbach (Addgene plasmid #71237; RRID:Addgene_71237), using annealed oligos and the BsmBI cloning site. Lentivirus for each guide was produced as described above and iPSCs were transfected with MOI 7.5 of LacZ or one of the *LINC00662*-targeting guide RNAs.

LINC00662 guide 1: CCCTGTGCGAGGTCTAACCC
LINC00662 guide 2: TGCTCCATAGGAGGACCCAT
LacZ: TGCGAATACGCCCACGCGAT

#### GFP+ cell isolation of transduced iPSCs

7 days after transduction, cells were detached with Accutase, resuspended in iPS media containing RY27632 (10 μM) and Draq7 (1:1000) and strained with a 70μm filter. Gating parameters were set by side and forward scatter to eliminate debris and aggregated cells. The GFP^+^ gates were set using untransduced iPSCs. The sorting gates were validated via reanalysis of sorted cells (> 99% purity cut-off). 300000 GFP^+^ /Draq7^-^ cells were collected per sample, spun down at 400 × g for 5 min, resuspended in iPS media containing RY27632 (10 μM) and expanded as described above and frozen down for further use. Guide efficiency was validated using standard quantitative real-time RT-PCR.

#### LINC00662-CRISPRi unguided neural organoids

*LINC00662*-CRISPRi organoids were generated as described above for human iPSC lines (see: Unguided neural organoid culture) with the exception that *LINC00662*-CRISPRi organoids were grown for 30 days. 2 independent organoid batches were generated. Day 15 and day 30 organoids were collected for downstream analyses. 5 day-15 and 3 day-30 organoids were collected and snap-frozen for each snRNA-seq replicate (n=2 per condition per batch). 5 day-15 and 3 day-30 organoids were collected for immunostainings (see: Immunohistochemistry) per condition per batch.

### Human tissue

Human fetal forebrain tissue was obtained from material available following elective termination of pregnancy at the University Hospital in Malmö, Sweden, in accordance with the national ethical permit (Dnr 2022-05051-01).

## Data availability

All data needed to evaluate the conclusions in the paper are present in the paper and/or the Supplementary Materials. The sequencing data presented in this study has been deposited at ENA, accession no: PRJEB104347. Proteomic data has been deposited to the ProteomeXchange Consortium via the PRIDE partner repository with the identifier PXD071764. Original code has been deposited at GitHub and is publicly available at github.com/Molecular-Neurogenetics/LINC00662_PTERV_Gerdes2025.

## Acknowledgements

We would like to thank D. Trono, F. H. Gage, W. Huttner, and S. Pääbo for comments and support. We thank U. Jarl and A. Hammarberg for technical assistance. We would like to acknowledge the support of the National Genomics Infrastructure (NGI) / Uppsala Genome Center for assisting in sequencing. We would also like to thank the Center for Translational Genomics (CTG) Lund University for providing expertise and service with sequencing and analysis.

## Supplementary Figures

**Figure S1.**
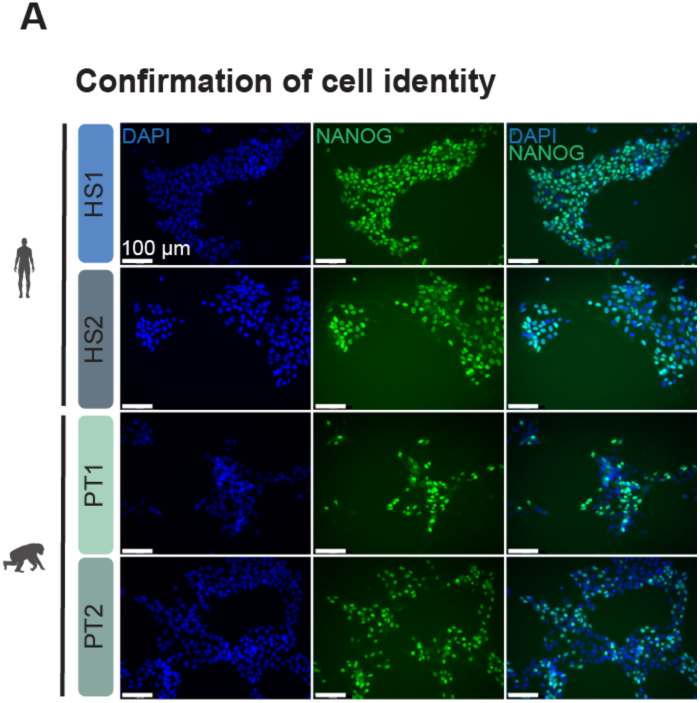
**A** Immunocytochemistry of human and chimpanzee iPSCs. Pluripotent cells stained with NANOG (green), nuclei stained with DAPI (blue). Scale bars = 100 μm.

**Figure S2.**
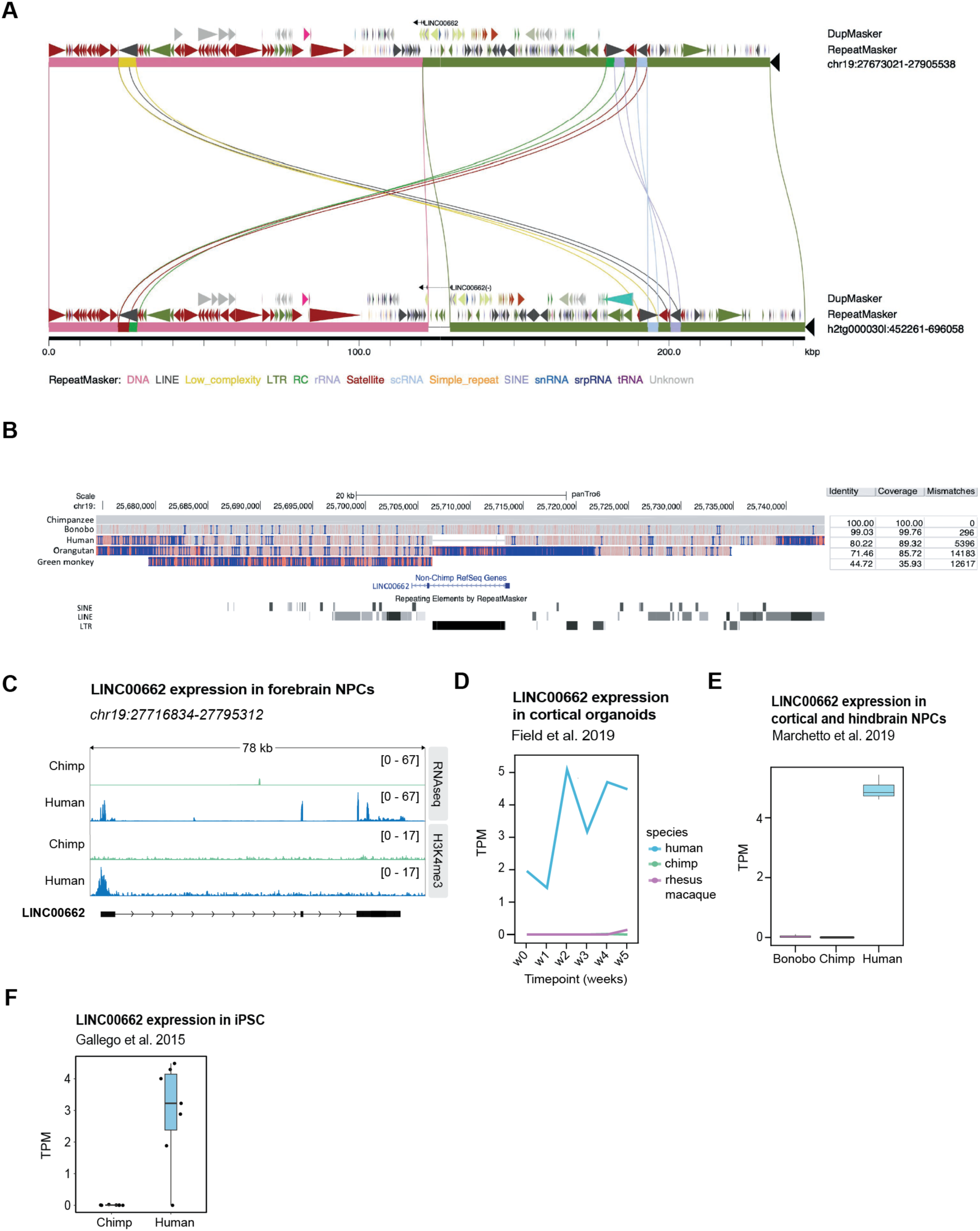
**A** Miropeats analysis reveals the PTERV1 insertion sequence in Clint_PTR (h2tg000030l; NCBI: PRJNA941354). Lines indicate sequence homology between the human reference (GRCh38, top) and Clint_PTR sequences. Additional annotations include segmental duplication (DupMasker) and RepeatMasker tracks. **B** Genome browser snapshot of LINC00662 locus in Pantro6. Top left: Conservation scores of the region across human, chimpanzee, bonobo, orangutan and green monkey. Top right: identity, coverage score, and number of mismatches to PanTro6 for each species. Middle: Gene body of LINC00662. Bottom: RepeatMasker annotation. **C** Genome browser snapshot of LINC00662 locus in hg38. Top two tracks show RNA-seq data from chimp and human fbNPCs respectively. Bottom two tracks showing H3K4me3 CUT&RUN data of chimp and human fbNPCs respectively. **D** Transcripts per million (TPM) of LINC00662 in human (blue), chimp (green), and rhesus macaque (purple) cortical organoids as reported by Field et al. 2019. **E** TPM of LINC00662 in human (blue), chimp (green) and bonobo (purple) in cortical and hindbrain NPCs as reported by Marchetto et al. 2019. **F** TPM of LINC00662 in human (blue) and chimp (green) in iPSCs as reported by Gallego et al. 2015.

**Figure S3.**
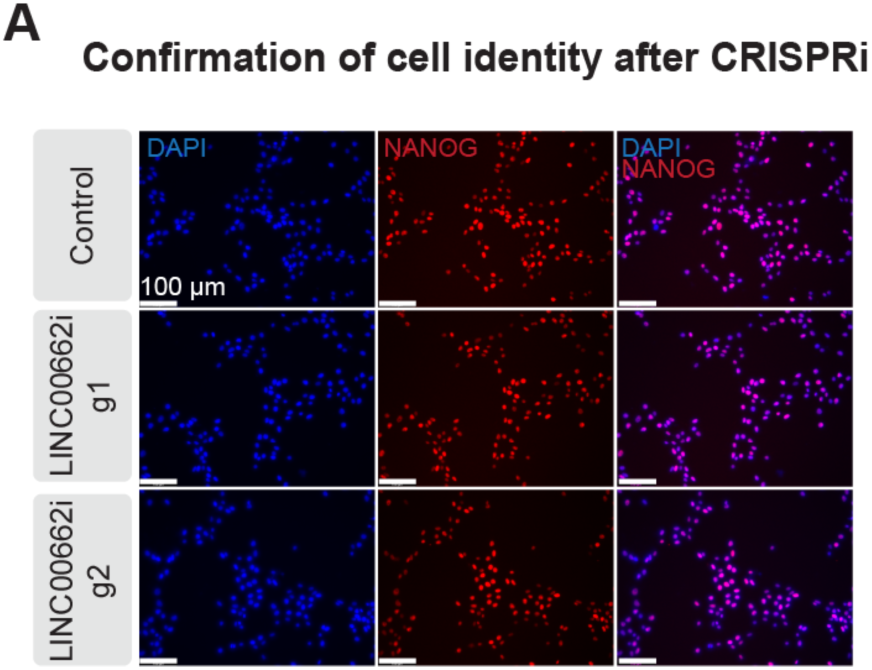
**A** Immunocytochemistry of *LINC00662*-CRISPRi iPSCs. Pluripotent cells stained with NANOG (red), nuclei stained with DAPI (blue). Scale bars = 100 μm.

